# A likelihood-free estimator of population structure bridging admixture models and principal components analysis

**DOI:** 10.1101/240812

**Authors:** Irineo Cabreros, John D. Storey

## Abstract

We introduce a simple and computationally efficient method for fitting the admixture model of genetic population structure, called `ALStructure`. The strategy of `ALStructure` is to first estimate the low-dimensional linear subspace of the population admixture components and then search for a model within this subspace that is consistent with the admixture model’s natural probabilistic constraints. Central to this strategy is the observation that all models belonging to this constrained space of solutions are risk-minimizing and have equal likelihood, rendering any additional optimization unnecessary. The low-dimensional linear subspace is estimated through a recently introduced principal components analysis method that is appropriate for genotype data, thereby providing a solution that has both principal components and probabilistic admixture interpretations. Our approach differs fundamentally from other existing methods for estimating admixture, which aim to fit the admixture model directly by searching for parameters that maximize the likelihood function or the posterior probability. We observe that `ALStructure` typically outperforms existing methods both in accuracy and computational speed under a wide array of simulated and real human genotype datasets. Throughout this work we emphasize that the admixture model is a special case of a much broader class of models for which algorithms similar to `ALStructure` may be successfully employed.

## 1 Introduction

Understanding structured genetic variation in human populations remains a foundational problem in modern genetics. Such an understanding allows researchers to correct for population structure in GWAS studies, enabling accurate disease-gene mapping (Knowler *et al*., 1988; Marchini *et al*., 2004; Song *et al*., 2015). Additionally, characterizing genetic variation is important for the study of human evolutionary history (Cavalli-Sforza *et al*., 1988; Esteban *et al*., 1998; Li *et al*., 2008).

To this end, much work has been done to develop methods to estimate what Alexander *et al*. (2009) term *global ancestry*. In the global ancestry framework, the goal is to simultaneously estimate two quantities:

(i) the allele frequencies of ancestral populations
(ii) the admixture proportions of each modern individual

Many popular global ancestry estimation methods have been developed within a probabilistic framework. In these methods, which we will refer to as *likelihood-based* approaches, the strategy is to fit a probabilistic model to the observed genome-wide genotype data by either maximizing the likelihood function (Alexander *et al*., 2009; Tang *et al*., 2005) or the posterior probability (Gopalan *et al*., 2016; Pritchard *et al*., 2000; Raj *et al*., 2014). The probabilistic model fit in each of these cases is the *admixture model*, described in detail in Section 2.1, in which the global ancestry quantities (i) and (ii) are explicit parameters to be estimated.

A related line of work relies on principal components analysis (PCA) and other eigen-decomposition methods, rather than directly fitting probabilistic models; as such, we will refer to them collectively as *PCA-based* approaches. These methods find many of the same applications as global ancestry estimates while obviating a direct computation of global ancestry itself. For example, the `EIGENSTRAT` method of Patterson *et al*. (2006) and Price *et al*. (2006) uses the principal components of observed data to correct for population stratification in GWAS, avoiding altogether the estimation of admixture proportions or ancestral allele frequencies. Similarly, Hao *et al*. (2016) observe that many important applications of global ancestry really only require *individual-specific allele frequencies*. In a sense, individual-specific allele frequencies are simpler than global ancestry; while global ancestry specifies all of the individual-specific allele frequencies, the converse is not true. Therefore, Hao *et al*. (2016) introduce a simple truncated-PCA method that accurately and efficiently estimates individual-specific allele frequencies alone.

Both likelihood-based and PCA-based approaches have distinct merits and drawbacks. The PCA-based methods are computationally efficient and accurate in practice. It is shown, for instance, that the individual-specific allele frequencies obtained by truncated-PCA are empirically more accurate than those obtained by likelihood-based methods (Hao *et al*., 2016). Another attractive feature of PCA-based methods is that they make minimal assumptions about the underlying data-generative model. However, as mentioned before, PCA-based methods do not provide the full global ancestry estimates that their corresponding likelihood-based methods do. Most notably, they do not provide direct estimates of admixture proportions, which are often of primary interest in some applications. Additionally, PCA-based methods often have weaker statistical justifications, as they are typically not based on a probabilistic model^1^.

Recognizing the relative advantages of each approach, several researchers have attempted to bridge the gap between likelihood-based and PCA-based approaches. In spirit, this is also the approach that we take in the present work, and so we briefly review previous contributions to contextualize the advances made by our own method. Engelhardt and Stephens (2010) observed that fitting the admixture model was related to PCA in the sense that both could be posed as matrix factorization problems, which differ only in the constraints imposed on factors. They then introduced a third matrix factorization problem, called Sparse Factor Analysis (SFA), which encourages a sparsity through a particular prior. However, since SFA does not enforce the probabilistic constraints of the admixture model (nor the orthogonality constraints of PCA), its output cannot be directly interpreted as an estimate of global ancestry. Lawson *et al*. (2012) provided further insight into the mathematical relationship between admixture models and PCA and introduced a method for the analysis of phased haplotype data. This method, called `fineSTRUCTURE`, fits a version of the admixture model in which each observed individual belongs to a single (rather than admixed) population. Zheng and Weir (2016) introduced a method called `EIGMIX` that leverages PCA to infer admixture proportions from unphased genotype data. While `EIGMIX` allows individual genomes to be derived from a mixture of multiple ancestral populations (unlike `fineSTRUCTURE`), it requires a set of sampled individuals known to be derived from single ancestral populations. A related line of work uses PCA-based approaches to fit models of *local ancestry*, in which inferences about the ancestry of individual genetic loci are desired (for example, Brisbin *et al*. (2012)).

While the aforementioned literature illustrates that PCA can be leveraged to provide information about population structure, each approach falls short of providing complete estimates of global ancestry under the general admixture model. The method which we introduce in the present work, called `ALStructure`, does precisely this. `ALStructure` requires no additional assumptions (such as the existence of unadmixed individuals in Zheng and Weir (2016)), no specialized input (such as the unphased haplotypes of Lawson *et al*. (2012)), and provides direct estimates of admixture proportions (unlike Engelhardt and Stephens (2010)). As such, `ALStructure` is the only existing PCA-based method that can provide a direct substitute to the most popular likelihood-based approaches. As an additional important advantage, the underlying mathematical theory which justifies `ALStructure` is sufficiently general so as to apply to a class of models that subsumes the admixture model. As such, we believe that imitable algorithms to `ALStructure` could be useful beyond the present genetics application.

The basic strategy of `ALStructure` is to eliminate the primary shortcomings of PCA-based methods while retaining their important advantages over likelihood-based methods. In particular we extend the approach taken in Hao *et al*. (2016) in two ways. First, we replace classical PCA with the closely related method of *Latent Subspace Estimation* (LSE) (Chen and Storey, 2015). In so doing, we will make mathematically rigorous the empirically effective truncated-PCA method of Hao *et al*. For estimating individual-specific allele frequencies. Second, we use the method of *alternating least squares* (ALS) to transform the individual-specific allele frequencies obtained via LSE into estimates of global ancestry.

We perform a body of simulations and analyze several globally and locally sampled human studies to demonstrate the performance of the proposed method, showing that `ALStructure` typically outperforms existing methods both in terms of accuracy and speed. We also discuss its implementation and the trade-offs between theoretical guarantees and run-time. We find that `ALStructure` is a computationally efficient and statistically accurate method for modeling admixture and decomposing systematic variation due to population structure.

The remainder of this paper is organized as follows. Section 2 introduces the admixture model and details the mathematical underpinnings of our approach. Section 3 summarizes the `ALStructure` algorithm. A reader primarily interested in a basic understanding of the operational procedure of `ALStructure` and its applications may proceed to Section 3 after reading Section 2.1, as the remainder of Section 2 is more technical in nature. Sections 4 and 5 assess the performance of `ALStructure` on a wide range of real simulated datasets.

## 2 Model and theory

In this section and the following we present the `ALStructure` method and detail some of its mathematical underpinnings. In Section 2.1, we define the *admixture model*: the underlying probabilistic model assumed by `ALStructure`. Section 2.2 describes the overall strategy of `ALStructure` as an optimality search subject to constraints rather than navigating a complex likelihood surface. Section 2.3 describes how the constraints can be used to estimate individual-specific allele frequencies. In Section 2.4 we present a mathematical result from Chen and Storey (2015) upon which the `ALStructure` algorithm heavily relies. Section 2.5 describes why estimating global ancestry, given the individual-specific allele frequencies, is equivalent to a constrained matrix factorization problem. An efficient algorithm based on the method of alternating least squares (ALS) is also provided in this section for performing the constrained matrix factorization. The complete `ALStructure` algorithm is then presented in Section 3.

Throughout this work, we adhere to the following notational convention: for a matrix ***A***, we denote the *i* row vector of ***A*** by ***a**_i•_*, the *j* column vector of ***A*** by ***α**_•j_*, and the (*i, j*) element of ***A*** by *a_ij_*.

### 2.1 The admixture model

The observed data ***X*** is an *m* × *n* matrix in which *m* (the number of SNPs) is typically much larger than *n* (the number of individuals). An element *x_ij_* of ***X*** takes values 0, 1, or 2 according to the number of reference alleles in the genotype at locus *i* for individual *j*.

`ALStructure` makes the assumption common to all likelihood-based methods that the data are generated from the *admixture model*. Under this model, the genotypes are generated independently according to *x_ij_*/*f_ij_* ~ Binomial(2, *f_ij_*), where ***F*** is an *m* × *n* matrix encoding all of the binomial parameters. Each element *f_ij_* is an *individual-specific allele frequency*: the frequency of allele *i* in individual *j*. ***F*** is further assumed to be of rank *d*, where *d* ≪ *n* ≪ *m*. *d* may be thought of as the number of ancestral populations from which the observed population is derived. ***F*** then admits a factorization ***F*** = ***PQ*** in which ***P*** and ***Q*** have the following properties:

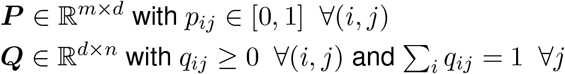

The matrices ***P*** and ***Q*** have the following interpretations: (i) each row ***p**_i•_* of ***P*** represents the frequencies of a single SNP for each of the *d* ancestral populations and (ii) each column ***q**_•j_* of ***Q*** represents the admixture proportions of a single individual. Together, ***P*** and ***Q*** encode the global ancestry parameters of the observed population; the goal of existing likelihood-based methods is to estimate these matrices. By contrast, the truncated-PCA method of Hao *et al*. (2016) is focused on estimating ***F*** and not its factors. Eq. 1 summarizes the admixture model.

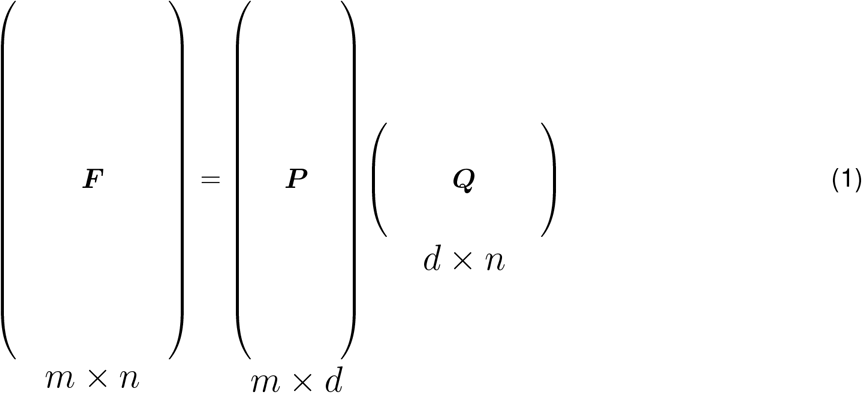

The model introduced in Pritchard, Stephens and Donnelly (2000), which we refer to as the *PSD model*, is an important special case of the admixture model. It additionally assumes the following prior distributions^2^ on ***P*** and ***Q***:

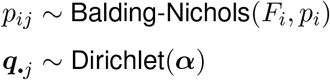

The Balding-Nichols distribution (Balding and Nichols, 1995) is a reparameterization of the Beta distribution in which *F_i_* is the *F_ST_* (Weir and Cockerham, 1984) at locus *i* and *p_i_* is the population minor allele frequency at locus *i*. Specifically, Balding-Nichols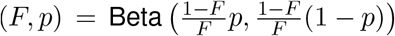. Existing Bayesian methods (Gopalan *et al*., 2016; Pritchard *et al*., 2000; Raj *et al*., 2014) fit the PSD model specifically, while existing maximum likelihood methods (Alexander *et al*., 2009; Tang *et al*., 2005) and `ALStructure` require only the admixture model assumptions.

Although we focus on fitting the admixture model in the present work, it is important note that the general strategy of the `ALStructure` algorithm is insensitive to the particular details of this model. The necessary features that the theoretical underpinnings of `ALStructure` require are: 1) higher moments of *x_ij_* are bounded, 2) ***F*** is low rank, and 3) *m* ≫ *n*.^3^ For example, an imitable algorithm could be applied to high dimensional data ***X*** where *x_ij_*|*f_ij_* ~ Poisson(*f_ij_*) and ***F*** is a low rank matrix whose factors ***P*** and ***Q*** potentially have natural constraints. Because of its generality, we hope that the approach of `ALStructure` will find useful application beyond the analysis of admixture.

### 2.2 Optimal model constraints

Most existing methods for fitting the admixture model employ various optimization techniques to search for the maximum likelihood parameters (Alexander *et al*., 2009; Pritchard *et al*., 2000) or the maximum a posteriori estimate (Gopalan *et al*., 2016; Raj *et al*., 2014). Our approach has a fundamentally different character: rather than searching through a rough likelihood landscape in pursuit of an optimal solution, `ALStructure` seeks a feasible solution to a set of optimal constraints. To be more specific, we begin with the observation that any solution satisfying a particular set of constraints is risk-minimizing among a class of unbiased estimators. Because any feasible solution is optimal, we can think of the constraints themselves as being optimal. Notably, the need to maximize likelihood is altogether obviated.

The challenge of this approach is twofold. First, the constraints themselves need to be estimated from the data: they are not directly observable. This is done through the method of Latent Subspace Estimation (LSE) detailed in Section 2.4. Second, feasible solutions to the estimated constraints will not typically exist. For this reason, we seek solutions that approximately satisfy the constraints, thereby converting a feasibility problem to a least squares optimization problem. This procedure is done through the method of Alternating Least Squares (ALS) and is detailed in Section 2.5. Throughout the remainder of the present subsection, we detail the constraints themselves.

There are several constraints that any reasonable estimate of the parameters of the admixture model must obey. The first is simply that the parameter estimates 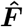, 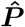, and 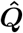 obey the relationship 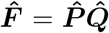. We will refer to this constraint as the *Equality* constraint. The second obvious requirement is that entries of matrices 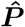 and 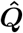 obey the probabilistic constraints of the admixture model:

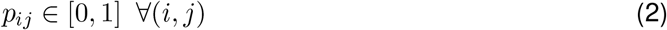

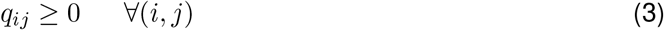

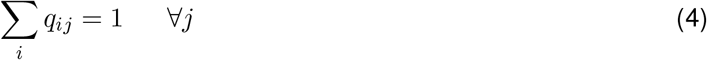

As we will encounter these constraints frequently, we refer to Eq. (2) as the “□” constraint, and Eq. (3) and (4) as the “Δ” constraint. This is simply because the constraints on ***P*** demarcate the boundaries of a *d*-dimensional unit cube (the generalization of a square) whereas the constraints on ***Q*** demarcate a d-dimensional simplex (the generalization of an equilateral triangle). Together we refer to the □ and Δ constraints as the *Boundary* constraints.

The final constraint we require is that the row vectors of 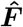 lie in the linear subspace spanned by the rows vectors of ***Q***. If we denote 〈***A***〉 to be the rowspace of a matrix ***A***, we can summarize this condition as:

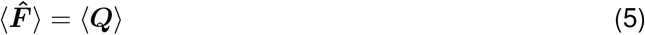

We will refer to Eq. 5 as the *LS* (linear subspace) constraint. The LS constraint is the only nontrivial constraint that `ALStructure` enforces. The fact that 〈***F***〉 = 〈***Q***〉 is a simple consequence of the linearity of the admixture model; indeed, all rows of ***F*** are linear combinations of rows of ***Q*** since ***F*** = ***PQ***. The LS constraint thus requires the same property for our estimate 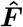. It is important to note that the LS constraint is not the same as requiring that 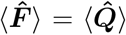: this is ensured by the Equality constraint. Rather, the LS constraint requires that the row vectors of 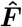 belong to the rowspace of the true ***Q*** matrix. The apparent challenge of enforcing the LS constraint is that *a priori*, one does not have access to 〈***Q***〉. However, `ALStructure` takes advantage of a recent result from Chen and Storey (2015) that 〈***Q***〉 can be consistently estimated directly from the data matrix ***X*** in the asymptotic regime of interest, when the number of SNPs m grows large. The result of Chen and Storey (2015) is in fact much more general than is needed in our setting and therefore will likely be useful in many other problems. Because of its importance to this work, we further discuss this result in the context of the admixture model in Section 2.4, and show that a modified PCA of ***X*** consistently recovers 〈***Q***〉.

### 2.3 Leveraging constraints to estimate 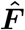

The key step in `ALStructure` is to note that enforcing the LS constraint provides us with an immediate estimate for ***F***. To motivate our estimator, first observe that the simple estimate 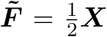 is in some sense a reasonable approximation of ***F***: it is unbiased since 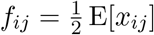 under the admixture model. However, this estimate leaves much to be desired — most importantly, the estimate 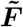 will in general be of full rank (*n*) rather than of low rank (*d*) and it will have a large variance. Assuming, for now, that we are provided with the true rowspace 〈***Q***〉 of ***F***, a natural thing to try is to project the rows of 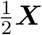 onto this linear subspace. Below we show that this estimator has some appealing properties.

Let us denote the the operator 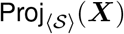 such that the rows of the matrix ***X*** are projected onto the linear subspace 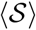.^4^ If we are given an orthonormal basis {*s_i_*} of the *d*-dimensional linear subspace 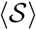, then:

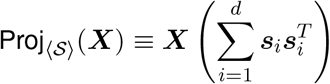

Lemma 1 below provides us a simple condition under which estimators of ***F*** formed by such projections are unbiased.

#### Lemma 1.

*For a rank d matrix **F** that admits a factorization **F** = **PQ** and a random matrix **X** such that* 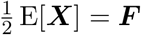, *any estimator of **F** of the form* 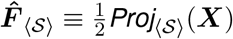 *is unbiased if and only if* 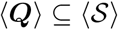.

Lemma 1 is proved in Appendix A.1. In particular we note that

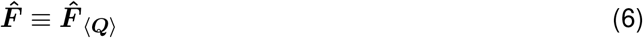

is unbiased.

In addition to being unbiased, the estimator 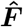 is optimal in the following sense. Among all unbiased estimators constructed by projecting ***X*** onto a linear subspace, 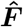 minimizes a matrix equivalent of the mean squared error.

#### Lemma 2.

*For a rank d matrix **F** that admits a factorization **F** = **PQ** and a random matrix X such that* 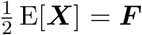, *the estimator* 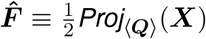 *is an unbiased estimator of **F** and has the smallest risk among all unbiased estimators of the form* 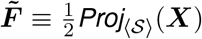. *We define the risk to be the expectation of the squared Frobenius norm:*

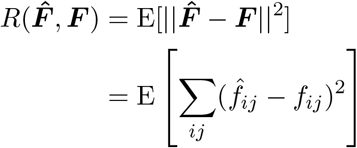

Lemma 2 is proved in detail in Appendix A.2, however the basic intuition is straightforward. Projecting ***X*** onto a linear space 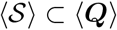 is biased (by Lemma 1). While projecting ***X*** onto a space 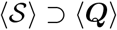 will result in an unbiased estimate of ***F*** (again, by Lemma 1), dimensions orthogonal to 〈***Q***〉 fit noise, increasing the variance (and therefore the mean squared error) of the estimate.

We note that this strategy is related to the strategy taken in Hao *et al*. (2016) in which ***F*** was estimated by projecting 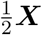 onto the space spanned by the first *d* principal components. In that work, it was observed that this simple strategy of estimating ***F*** typically outperformed existing methods. We will see in Section 2.4 that the space spanned by the first *d* principal components is a good estimator for 〈***Q***〉 itself, but it can be improved practically and with theoretical guarantees by performing a modified PCA. Therefore, Lemma 2 provides a theoretical justification for the empirically accurate method put forward in Hao *et al*. (2016).

### 2.4 Latent subspace estimation

We have shown that the linear subspace 〈***Q***〉 can be leveraged to provide a desirable estimate of ***F***. However, as 〈***Q***〉 is a linear subspace spanned by latent variables, it is not directly observable and must be estimated. Here we show how a general technique developed in Chen and Storey (2015), which we will refer to as *Latent Subspace Estimation* (LSE), can be used to compute a consistent estimate of 〈***Q***〉 from the observed data ***X***.

LSE is closely related to PCA, a popular technique that identifies linear combinations of variables that sequentially maximize variance explained in the data (Jolliffe, 2002). As PCA is commonly used to find low-dimensional structure in high-dimensional data, a natural approach to estimating 〈***Q***〉 would be to employ SNP-wise PCA. More specifically, we might consider the linear space spanned by the first few eigenvectors of the *n* × *n* sample covariance matrix, 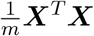 as an estimate of 〈***Q***〉.

The LSE-based estimate of 〈***Q***〉 almost exactly matches this PCA-based intuition. The only difference is that LSE accounts for the heteroscedastic nature of the admixture model, as detailed in Chen and Storey (2015). LSE has the theoretical advantage of asymptotically capturing 〈***Q***〉 in the highdimensional setting (i.e. as *m* → ∞). This is what is expressed in below in Theorem 1 below: a special case of a more general theorem from Chen and Storey (2015), rewritten here for the special case of the admixture model).

#### Theorem 1

(Chen and Storey (2015)). *Let us define* 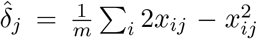 *and let **D** be the diagonal matrix with jth entry equal to* 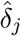. *The d eigenvectors* {*v*_1_,…, *v_d_*} *corresponding to the top d eigenvalues of the matrix* 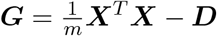 *span the latent subspace* 〈***Q***〉 *in the sense that*

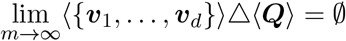

*with probability 1, where* Δ *denotes the symmetric set difference. Further, the smallest n* – *d eigenvalues of **G** converge to 0 with probability 1*.

Theorem 1 provides us with a simple procedure for estimating 〈***Q***〉 directly from data. One first computes 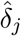 and constructs the *n* × *n* matrix ***D***. Next, an eigendecomposition of the adjusted covariance estimate 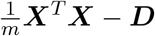 is computed. Finally, we estimate 〈***Q***〉 as

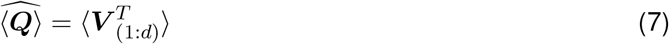

where ***V***_1:*d*_ are the first *d* columns from the singular value decomposition of ***G***.

We stress that the general form of Theorem 1 from Chen and Storey (2015) makes LSE applicable to a vast array of models beyond factor models and the admixture model discussed here. As a further benefit to the LSE methodology, it is both easy to implement and computationally appealing. The entire computation of 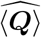 requires a single eigen-decomposition of an *n* × *n* matrix where the accuracy depends only on large *m*.

### 2.5 Leveraging constraints to estimate *P* and *Q*

Now that we have a method for obtaining the estimate 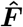 by leveraging the LS constraint, what remains is to find estimates for ***P*** and ***Q***. Since the estimate 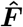 has several appealing properties, as outlined in Section 2.3, the approach of `ALStructure` is simply to keep 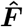 fixed and seek matrices 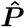 and 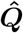 that obey the Equality and Boundary constraints of the admixture model. Below we discuss some of the general properties of this approach: namely the question of existence and uniqueness of solutions. We will briefly discuss the general problem of non-identifiability in the admixture model and provide simple and interpretable conditions under which the admixture model is identifiable. Finally, we will provide simple algorithms for computing 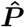 and 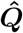 from 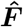 based on the method of Alternating Least Squares (ALS).

#### Existence, uniqueness, and anchor conditions

First we develop some terminology. We will say that an *m* × *n* matrix ***A** admits an admixture-factorization* if the following feasibility problem has a solution:

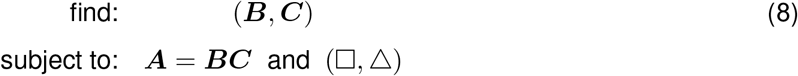

In words, the feasibility problem in Eq. 8 simply seeks a factorization of ***A*** that obeys the Equality and Boundary constraints from Section 2.2 imposed by the admixture model. The smallest integer *d* for which (***B, C***) is a solution to Eq. 8 with ***B*** an *m* × *d* matrix and ***C*** a *d* × *n* matrix is the *admixture-rank* of ***A***, which we denote rank_ADM_(***A***). By seeking a rank *d* admixture-factorization of 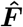, `ALStructure` converts a problem of high-dimensional statistical inference to a matrix factorization problem.

This simple approach has two apparent shortcomings:

i. A rank *d* admixture-factorization of 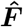 may not exist.
ii. If a valid factorization exists, it will not be unique.

Item (i) is a technical problem; though ***F*** admits a rank *d* admixture factorization by assumption, the same is not true for 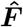 in general. Even though the rank of 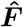 is *d* by construction, 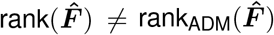 in general. `ALStructure` avoids this problem changing the feasibility problem expressed in Eq. 8 to the following optimization problem:

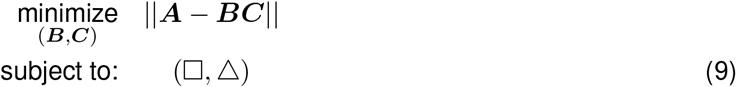

It is important to note that (ii) is not a problem unique to `ALStructure`, but is a fundamental limitation for any maximum likelihood (ML) method as well. This is because the likelihood function depends on 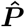 and 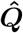 only through their product 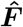; more formally, the admixture model is non-identifiable. One unavoidable source of non-identifiability is that any solution 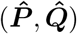 to the matrix factorization problem in Eq. 8 will remain a valid solution after applying a permutation to the columns of 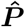 and the rows of 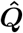. A natural question to ask is: “When is there a unique factorization 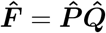 up to permutations?”

Two important types of sufficient conditions under which unique factorizations exist up to permutations are (i) *anchor SNPs* and (ii) *anchor individuals*. We note that the concept of anchors has been previously employed in the field of topic modeling, where *anchor words* are of interest (Arora *et al*., 2013). We define an anchor SNP as one that is fixed in all ancestral populations except one. The anchor SNPs condition is then satisfied if each of the *d* ancestral populations has at least one corresponding anchor SNP. Analogously we define an anchor individual as one whose entire genome is inherited from a single ancestral population. The anchor individuals condition then satisfied if each of the *d* ancestral populations has at least one corresponding anchor individual. The assumption of anchor individuals is equivalent to the assumption of “surrogate ancestral samples “ required by the EIGMIX method of Zheng and Weir (2016). The fact that either a set of *d* anchor SNPs or *d* anchor individuals makes the admixture model identifiable up to permutations follows from a simple argument found in Appendix A.3. For the special case of *d* = 3, Fig. 1 graphically displays the anchor conditions. It is important to remember that `ALStructure` does not *require* anchors to function. Rather, anchors provide interpretable conditions under which solutions provided by `ALStructure`, or any likelihood-based method, can be meaningfully compared to the underlying truth.

**Figure 1:**
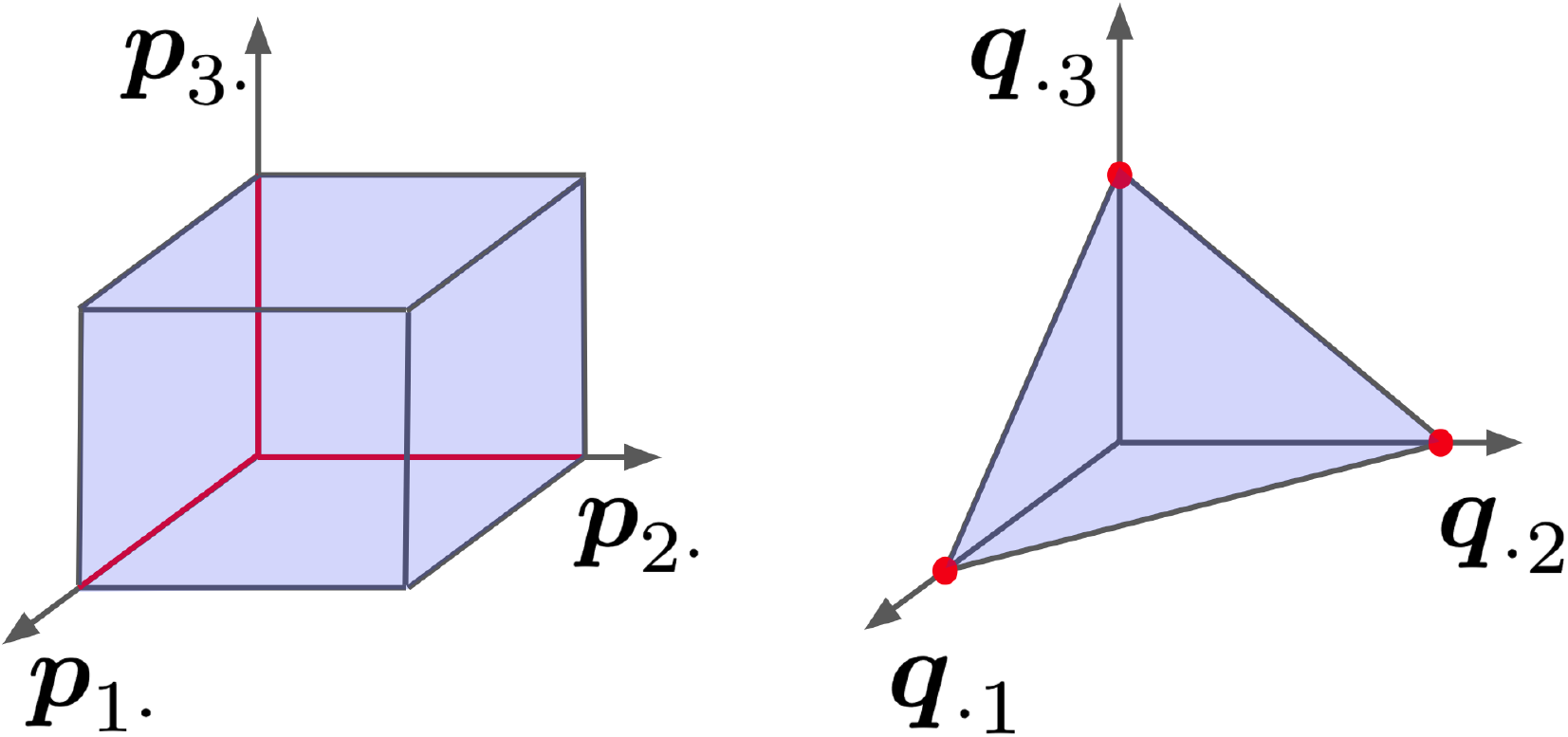
Summary of sufficient conditions for a factorization ***F*** = ***PQ*** to be unique for *d* = 3. Axes represent the components of the row vectors of ***P*** and the column vectors of ***Q*** respectively. (left) Anchor SNPs: there is at least one row of ***P*** on each on each of the red lines. (right) Anchor genotypes: there is at least one column of ***Q*** on each of the red dots.

The anchor SNP and anchor individual conditions are not necessarily the only sufficient conditions for ensuring identifiability of the admixture model and indeed to the best or our knowledge, there is not currently a complete characterization of conditions for which the admixture model is identifiable. We regard this as an important open problem. In practice, `ALStructure` is capable of retrieving solutions remarkably close to the underlying truth even in simulated scenarios far from satisfying the anchor conditions, including conditions which are challenging for existing methods.

#### Computation

Here we present two simple algorithms for solving the optimization problem:

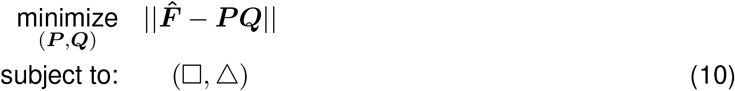

The first algorithm, which we call `cALS` (constrained Alternating Least Squares), has the advantage that it is guaranteed to converge to a stationary point of the nonconvex objective function in (10). While a stationary point will not generally correspond to a globally optimal solution, global optimization is seldom possible for nonconvex problems.

Although theoretically appealing, this algorithm relies on solving many constrained quadratic programming problems and is consequently potentially slow. To overcome this problem, we introduce a second algorithm called `tALS` (truncated Alternating Least Squares), which simply ignores the problematic quadratic constraints in `cALS`. Though lacking a theoretical guarantee of convergence, the increase in speed is significant and the outputs of the two algorithms are often practically indistinguishable.

We note that the general method of alternating least squares is not novel. In particular, previous work has developed alternating least squares methods for the the problem of nonnegative matrix factorization (NNMF) (Lee and Sebastian, 1999; Paatero and Tapper, 1994). In NNMF, one seeks a low-rank factorization ***A*** = ***BC*** in which all elements of the factors ***B*** and ***C*** are nonnegative. Algorithms analogous to `cALS` and `tALS`, but with nonnegetivity constraints rather than the □ and Δ constraints, have previously been considered (Berry *et al*., 2007; Cichocki *et al*., 2007; Gillis and Glineur, 2012; Kim *et al*., 2014).

##### An algorithm with provable convergence

While problem (10) is nonconvex as stated, the following two subproblems are convex:

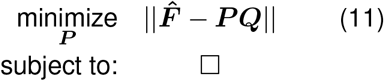

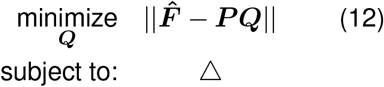

That (11) and (12) are convex problems is clear; norms are always convex functions and □ and Δ are convex constraints. In particular (11) and (12) are both members of the well-studied class of Quadratic Programs (QP) for which many efficient algorithms exist (Boyd and Vandenberghe, 2009). We propose the following procedure for factoring 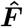, which we call *Constrained ALS Algorithm*.

###### Algorithm 1 Constrained ALS Algorithm

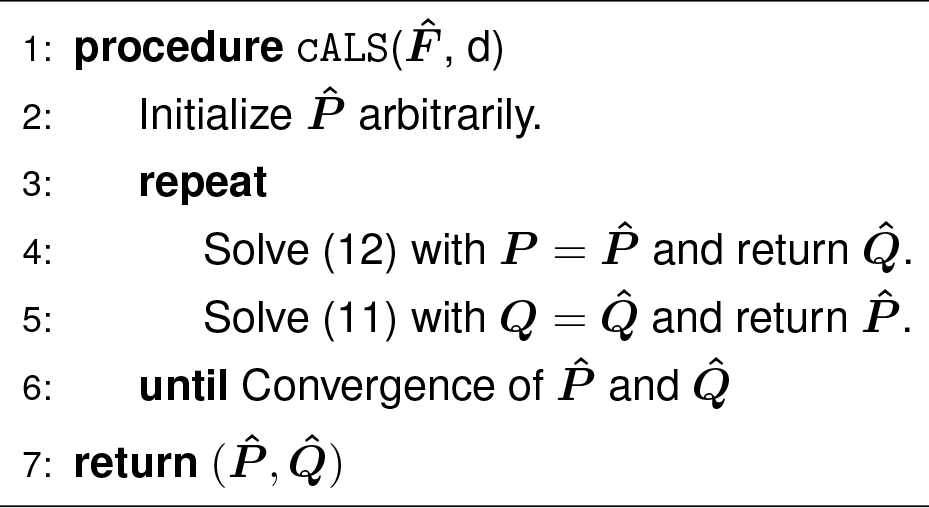

Despite the original problem being nonconvex, Algorithm 1 is guaranteed to converge to a stationary point of the objective function in (10) as a result of the following theorem from Grippo and Sciandrone (2000).

###### Theorem 2.

*For the two block problem*,

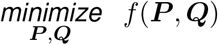

*if* {***P**_i_*} and {***Q**_i_*} *are a sequence of optimal solutions to the two subproblems:*

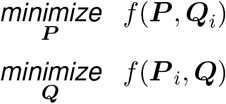

*then any limit point* (***P, Q***) *will be a stationary point of the original problem*.^5^

##### An efficient heuristic algorithm

If we remove all constraints on ***P*** and ***Q*** from Eq. 11 and 12, the resulting optimization problems are simple linear least squares (LS).

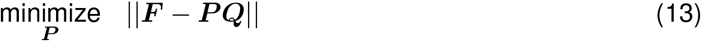

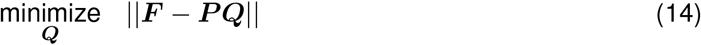

Our algorithm proceeds by alternating between solving the unconstrained LS problems (13) and (14). After each step, the optimal solution will not necessarily obey the constraints of problem (10). To keep our algorithm from converging on an infeasible point, we truncate the solution to force it into the feasible set. More precisely, each element of the solution ***P***\* to (13) is truncated to satisfy □ and each column of the solution ***Q***\* to (14) is projected to the closest point on the simplex defined by the Δ constraint. Simplex-projection is nontrivial, however it is a well-studied optimization problem. Here we use a particularly simple and fast algorithm from Chen and Ye (2011). This algorithm, which we call the *Truncated ALS Algorithm*, is detailed in Algorithm 2.

###### Algorithm 2 Truncated ALS Algorithm

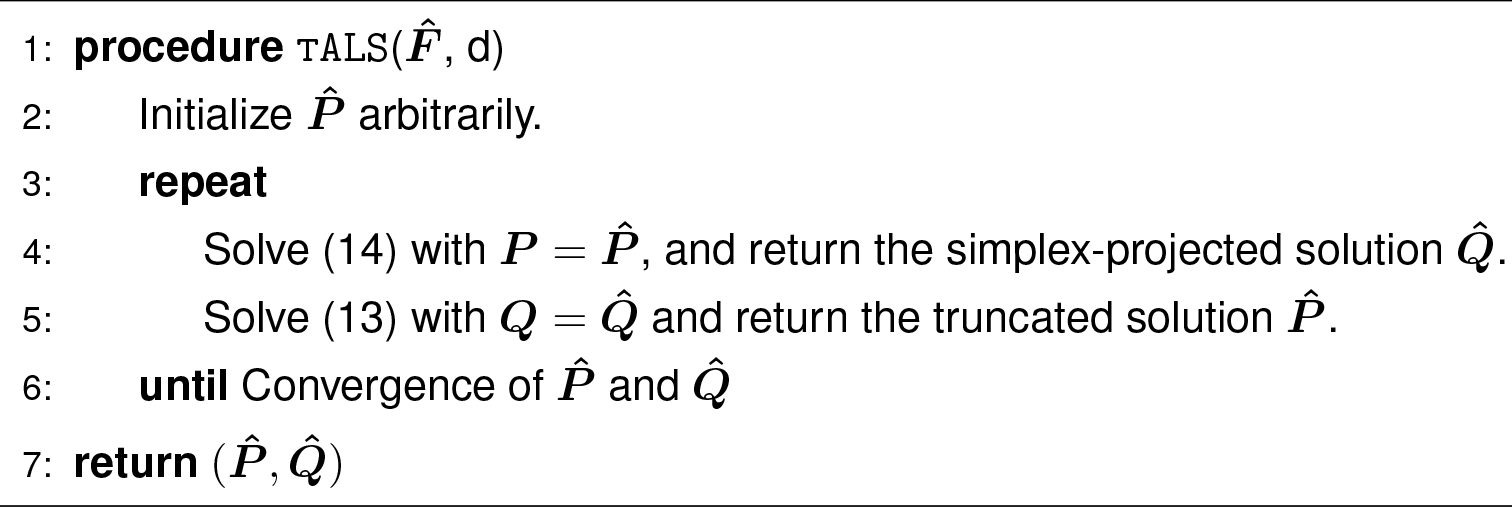

##### An example dataset

Figure 2 displays the output of `cALS` and `tALS` on a dataset from the PSD model with the parameters: *m* = 100 000, *n* = 500, *k* = 3, *α* = (0.1,0.1,0.1). As can be seen, the output fits for ***Q*** provided by `cALS` and `tALS` are practically indistinguishable to the eye and are both excellent approximations of the ground truth. The `cALS` algorithm performed slightly better than the `tALS` algorithm (8.5 × 10^−3^ and 8.7 × 10^−3^ RMSE, respectively). However, `cALS` took 3.5 hours to complete while `tALS` terminated in under 1.5 minutes. Because of the significant gains in efficiency, we use `tALS` exclusively throughout the remainder of this paper. The analyst who requires theoretical guarantees can, of course, use the `cALS` algorithm instead. Appendix B provides a more detailed comparison between the `tALS` and `cALS` algorithms on simulated data.

**Figure 2:**
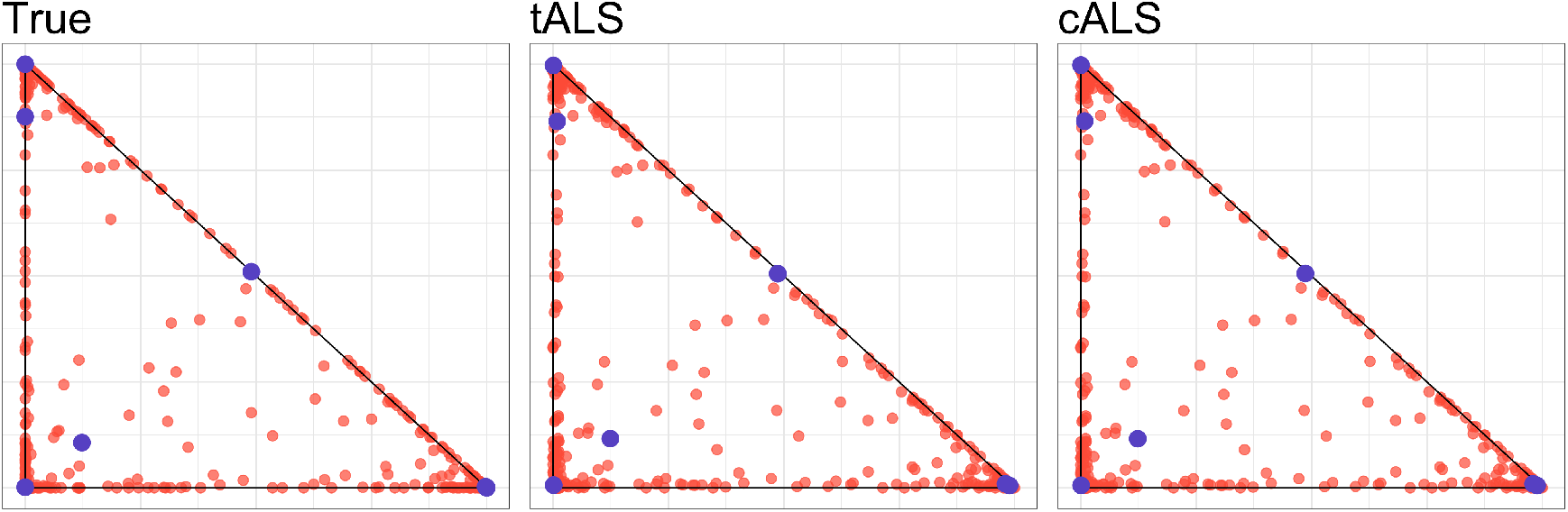
Biplots of the first two rows of ***Q*** (left), 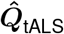 (middle) and 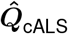 (right). Blue points are provided as a visual aid and delineate a common subset of individuals.

## 3 The ALStructure algorithm

In this section we briefly outline the entire `ALStructure` method whose components were motivated in depth in Section 2. In order to fit the admixture model, we obtain estimates 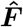, 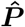, and 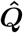 from the SNP matrix ***X*** through the following three part procedure:

(i) Estimate the linear subspace 〈***Q***〉 from the data ***X***.
(ii) Project 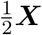 onto the estimate 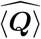 to obtain an estimate of ***F***.
(iii) Factor the estimate 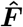 subject to the Equality and Boundary constriants to obtain 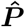 and 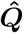.

For convenience, we detail the entire `ALStructure` algorithm in Algorithm 3 and annotate each of the three steps described above.^6^

We emphasize here that `ALStructure`’s estimate 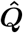 is ultimately derived from the LSE-based estimate of the latent subspace 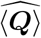. As the method of LSE is closely linked to PCA, we consider `ALStructure` to be a unification of PCA-based and likelihood-based approaches.

Perhaps the most striking feature of Algorithm 3 is its brevity. One advantage of this simplicity is its ease of implementation. Although Algorithm 3 has been implemented in the `R` package `ALStructure`, it can clearly be reimplemented in any language quite easily. Equally important is that all of the operations in Algorithm 3 are standard. The only two computationally expensive components are (i) a single eigen-decomposition (line 6) if *n* is large and (ii) QR decompositions to find linear least squares (LLS) solutions in the `tALS` algorithm. Both of these problems have a rich history and consequently have many efficient algorithms. It is likely that the `ALStructure` implementation of Algorithm 3 can be significantly sped up by utilizing approximate or randomized algorithms for the eigen-decomposition and/or LLS computations. In its current form, `ALStructure` simply uses the base `R` functions `eigen`() and `solve`() for the eigen-decomposition and LLS computations, respectively. Despite this, the current implementation of `ALStructure` is typically faster than existing algorithms as can be seen in Sections 4 and 5 below.

### Algorithm 3 ALStructure

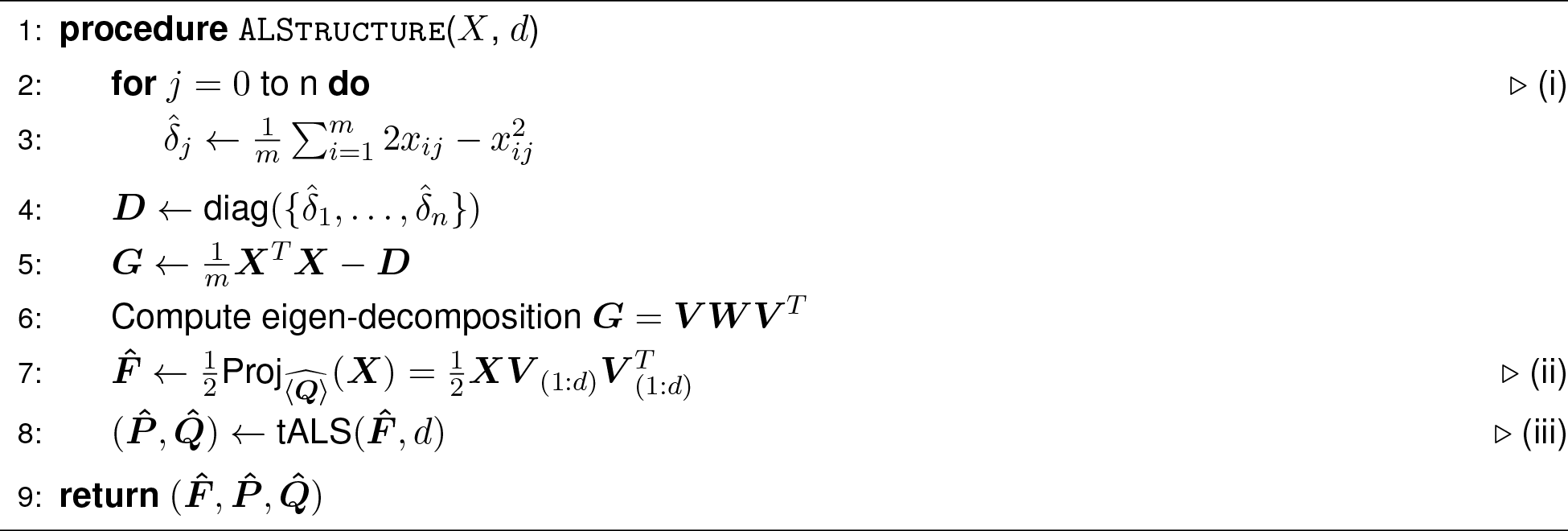

The `ALStructure` method is a nonparametric estimator in the following ways. It makes no assumptions about the probability distributions of ***P*** or ***Q***. Any random variable taking values in {0, 1} is by necessity Bernoulli. In this vein, the assumption that *x_ij_* ~ Binomial(2, *f_ij_*) is not a parametric assumption *per se*, but rather an assumption about independence of alleles. Finally, the likelihood function is not utilized in estimating ***P*** and ***Q***, making `ALStructure` likelihood-free.

For choosing the dimensionality of the model *d*, we recommend utilizing the recently proposed structural Hardy-Weinberg equilibrium (sHWE) test (Hao and Storey, 2017). This test can perform a genome-wide goodness of fit test to the assumptions made in the admixture model over a range of *d*. It then identifies the minimal value of *d* that obtains the optimal goodness of fit. There are other ways to choose *d*, by using the theory and methods in Chen and Storey (2015) or by using other recent proposals (Hao *et al*., 2016; Patterson *et al*., 2006).

## 4 Results from simulated data

### 4.1 Simulated data sets

In this section we compare the performance of `ALStructure` to three existing methods for global ancestry `estimation, Admixture, fastSTRUCTURE` and `terastructure`. `Admixture`, developed by Alexander *et al*. (2009), is a popular algorithm which takes a maximum likelihood approach to fit the admixture model. Both `fastSTRUCTURE` (Raj *et al*., 2014) and `terastructure` (Gopalan *et al*., 2016) are Bayesian methods that fit the PSD model using variational Bayes approaches. We abbreviate these methods as ADX, FS, and TS in the figures. A comparison among these three methods appears in Gopalan *et al*. (2016), so we will focus on how they compare to `ALStructure`.

To this end, we first tested all algorithms on a diverse array of simulated datasets. The bulk of our simulated data sets come from the classical PSD model (defined in Section 2.1) in which columns of ***Q*** are distributed according to the Dirichlet(*α*) distribution and the rows of ***P*** are drawn from the Balding-Nichols distribution. We varied the following parameters in our simulated datasets: *m, n, d*, and *α*. Of particular note is the variation of *α*. For this we used four *α*-prototypes: *α*_1_ = (10, 10, 10), *α*_2_ = (1, 1, 1), *α*_3_ = (0.1, 0.1, 0.1), and *α*_4_ = (10, 1, 0.1). These four prototypes were chosen because they represent four qualitatively different distributions on the Dirichlet simplex as shown in Fig. 3: *α*_1_ corresponds to points distributed near the center of the simplex, *α*_2_ corresponds to points distributed evenly across the simplex, *α*_3_ corresponds to points distributed along the edges of the simplex, and *α*_4_ corresponds to an asymmetric distribution in which points are concentrated around one of the corners of the simplex.

**Figure 3:**
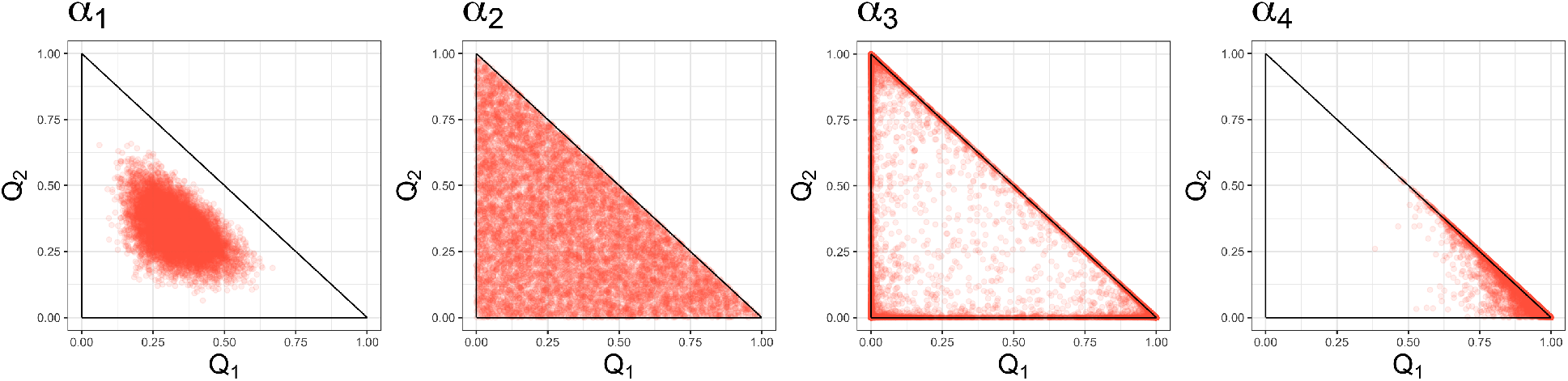
Examples of typical random samples from the four different *α*-prototypes. As can be seen, only *α*_2_ and *α*_3_ approximately obey the “anchor-individuals” condition.

When we produced datasets with *d* > 3, we extended the prototypes in the natural way; for example for *d* = 6, the *α*_4_ is becomes (10, 10, 1, 1, 0.1, 0.1). Table 1 lists all of the parameters we used to generate data under the Dirichlet model, for a total of 96 distinct combinations.

**Table 1:** Parameters of all simulated datasets

|  |  |
| --- | --- |
| $m$ | $10^5, 5 \times 10^5$ |
| $n$ | $5 \times 10^2, 10^3, 5 \times 10^3, 10^4$ |
| $d$ | 3, 6, 9 |
| α-prototypes | (10, 10, 10) |
|  | (1, 1, 1) |
|  | (0.1, 0.1, 0.1) |
|  | (10, 1, 0.1) |

The parameters of the Balding-Nichols distributions from which rows of the ***P*** matrix were drawn were taken from real data, following the same strategy taken in Gopalan *et al*. (2016). Specifically, *F_i_* and *p_i_* were estimated for each SNP in the Human Genome Diversity Project (HGDP) dataset (Cavalli-Sforza, 2005). Then for each simulated dataset, *m* random samples are taken (with replacement) from the HGDP parameter estimates.

In addition to simulating ***Q*** matrices from the classical Dirichlet(*α*) distribution with many different parameters *α*, we also simulated data from the spatial model of admixture developed in Ochoa and Storey (2016). We deliberately chose to study this model because it is ill-suited for `ALStructure`; while `ALStructure` relies on the estimation of the *d*-dimensional linear subspace 〈***Q***〉, the columns of ***Q*** produced under the spatial model lie on a one-dimensional curve within 〈***Q***〉. Despite this fundamentally challenging scenario, we see that `ALStructure` is often capable of recovering an accurate approximation.

### 4.2 Results from the PSD model

In order to give a representative picture of the relative performance of `ALStructure` against existing algorithms, we first plot the fits of all of the algorithms for two particular data sets out of the total 96 model data sets: (i) the data set in which `ALStructure` performs the best and (ii) the data set in which `ALStructure` performs the worst, according to mean absolute error (defined below).

In Fig. 4a, we see that all four algorithms perform very well for the data set in which `ALStructure` performs best, which comes from the *α*_3_-prototype. In Fig. 4b, the dataset was generated from the *α*_4_-prototype. We see that while `ALStructure` certainly deviates substantially from the truth, so does every algorithm. Both `fastSTRUCTURE` and `terastructure` provide results that are qualitatively very different from the truth; where `fastSTRUCTURE` compresses all columns of ***Q*** onto a single edge of the simplex, `terastructure` spreads them out through the interior of the simplex. Both `Admixture` and `ALStructure` provide solutions qualitatively similar to the truth. While the points in the `Admixture` solution extend much further along the edge of the simplex than the true model, the `ALStructure` solution spreads into the interior of the simplex more than the true model.

**Figure 4:**
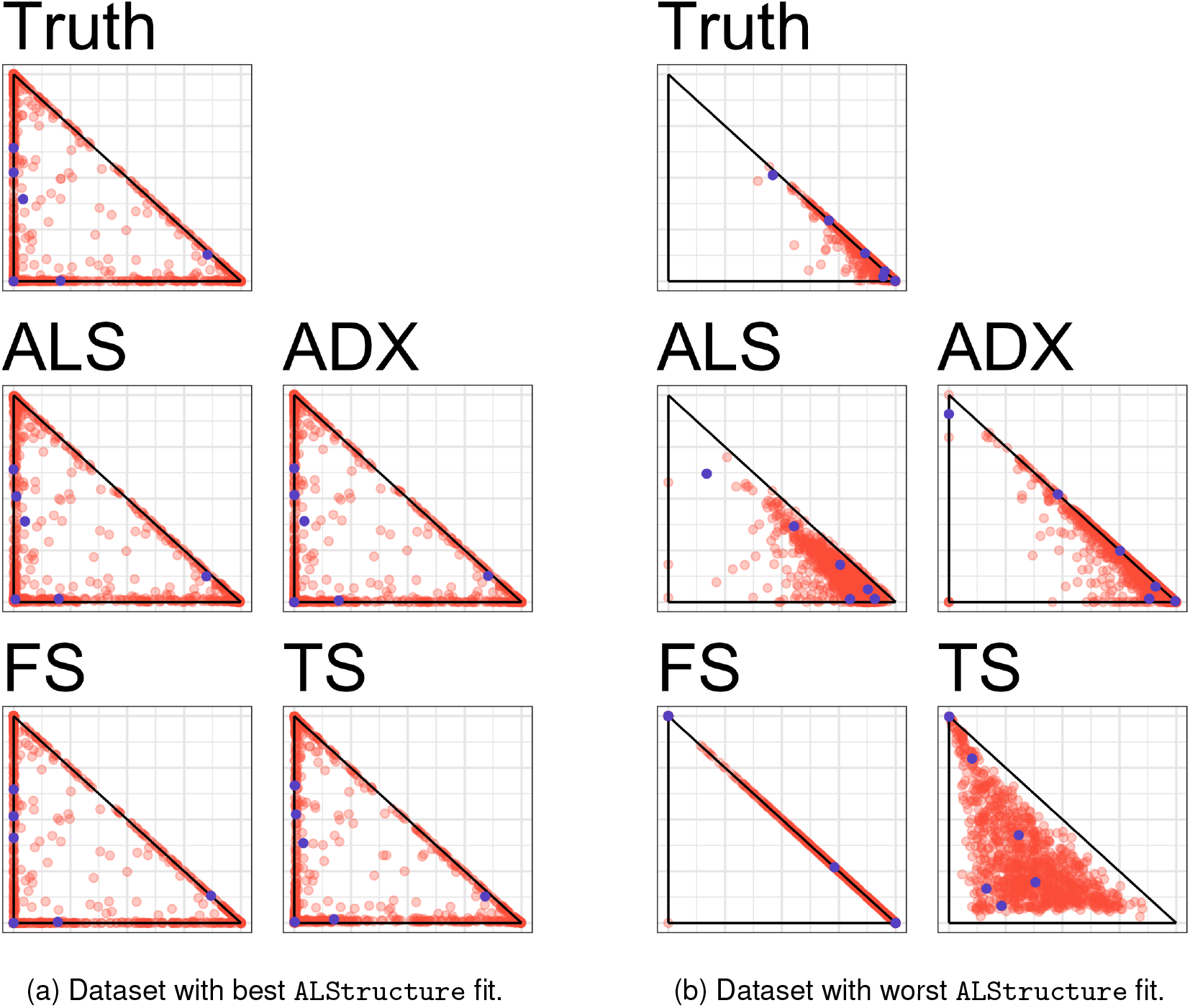
Model fits by `ALStructure, Admixture, fastSTRUCTURE`, and `terastructure` on the two particular simulated datasets. Each point represents a column of the ***Q*** matrix and is plotted by the first and second coordinates. Blue points are plotted as a visual aid and delineate a common subset of individuals.

Fig. 5 provides a comprehensive summary of the performance of `ALStructure` against the existing algorithms on all simulated datasets. The top panels of Fig. 5 summarize the accuracy of each of the four algorithms, according to two metrics: root mean squared error (RMSE) and mean absolute error (MAE).

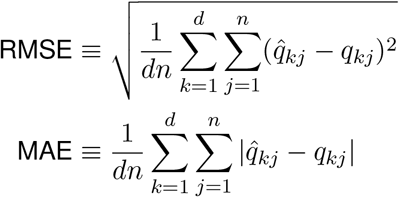

**Figure 5:**
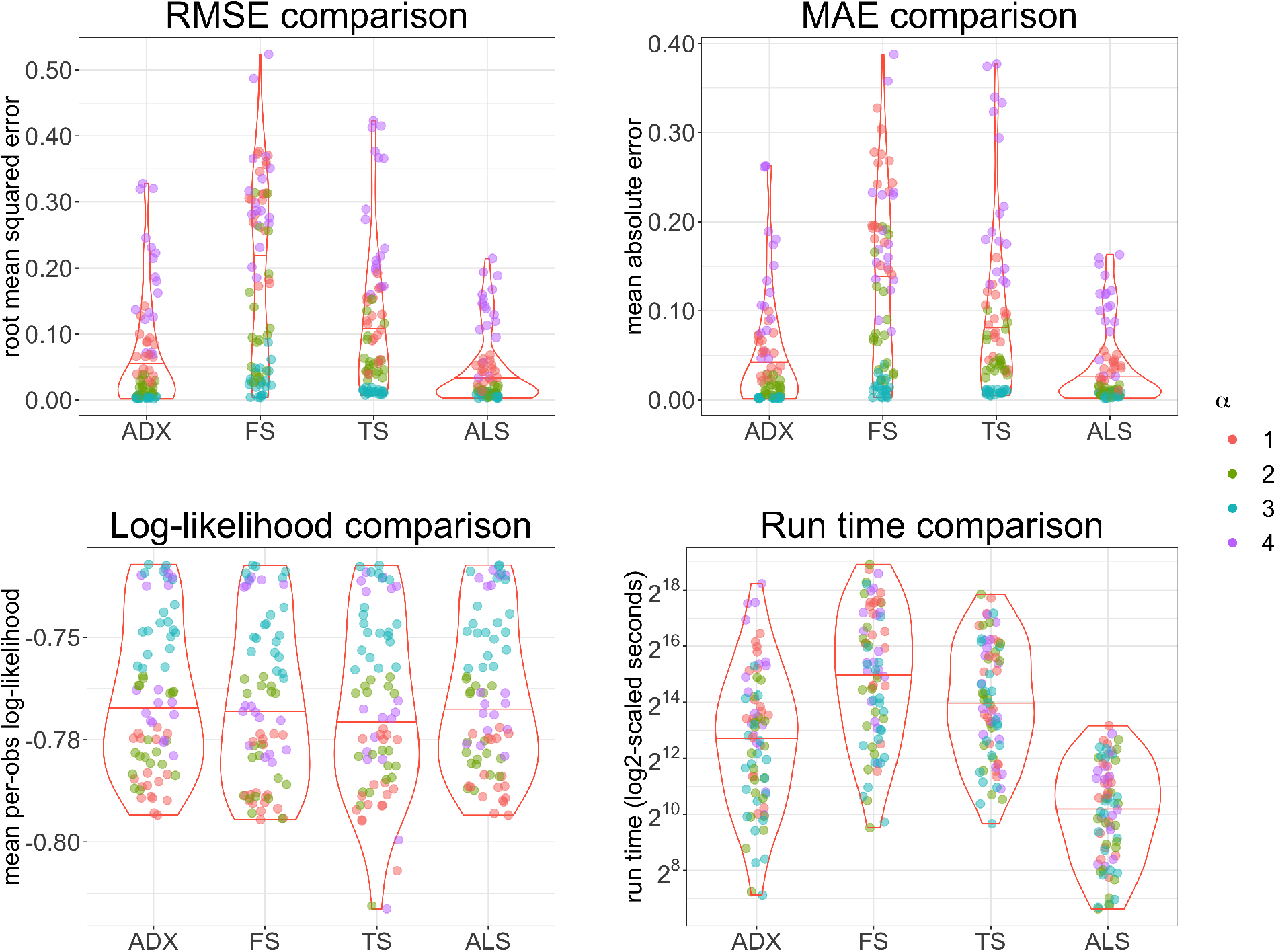
Summary of performance of `ALStructure` and existing algorithms. The points are colored by *α*-prototype.

The bottom left panel of Fig. 5 shows mean per-observation log-likelihood, 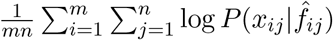, on all simulated data sets. (To obtain full data log-likelihoods, multiple these numbers by *mn*.) It is interesting to note that `ALStructure` performs comparably to other methods from the likelihood perspective despite the fact that it is the only method which does not explicitly utilize the likelihood function. However, we emphasize that likelihood is an imperfect metric of model fit for two reasons. First, because of the highly non-identifiable nature of the admixture model as discussed in Section 2.5, many models are equivalent from the likelihood perspective. Therefore, if one is primarily concerned about the accuracy of admixture estimates, the RMSE or MAE metrics may be more suitable. Second, in high-dimensional models it has been demonstrated that high likelihood may yield far inferior estimates (Efron, 2013). Starting with Stein’s Paradox (Stein, 1956), it has been shown in many settings that the maximum likelihood estimator for several parameters may be uniformly worse in accuracy than methods that leverage shared information in the data.

The bottom right panel of Fig. 5 shows the distributions of run times for each algorithm on all modeled datasets. Due to the size of the simulated datasets and our computational constraints, each algorithm did not terminate on each of the 96 datasets. In Fig. 5, we plot only the datasets for which all four algorithms successfully terminated. See Appendix C for more details. It is clear that `ALStructure` is competitive with respect to both model fit and time. `ALStructure` outperforms all methods according to both RMSE and MAE. With respect to time, `ALStructure` is clearly favored (one should note that the *y*-axis is on the `log` scale).

### 4.3 Results from the spatial model

As a challenge to `ALStructure`, we simulate data from a model developed in Ochoa and Storey (2016), which we will refer to as the *spatial model*. This model mimics an admixed population that was generated by a process of diffusion in a one-dimensional environment. There are *d* unmixed ancestral populations equally spaced at positions {*x*_0_, *x*_0_ +1, …, *x*_0_ + *d* − 1} on an infinite line. If all populations begin to diffuse at time *t* = 0 at the same diffusive rate, then population *i* will be distributed as a Gaussian with mean *μ_i_* = *x*_0_ + *i* − 1 and standard deviation *σ*. Therefore, under the Spatial model, an individual sampled from position *x* will have admixture proportions:

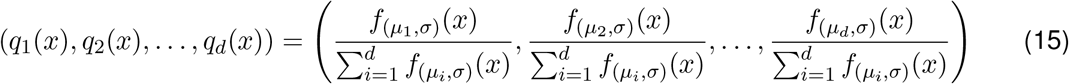

where *f*_(*μ σ*)_ denotes the Gaussian distribution with parameters (*μ, σ*).

Although this is just a special case of the admixture model, one would expect the spatial model to be particularly challenging for `ALStructure` because the admixture proportions belong to a onedimensional curve parameterized by *x*, and `ALStructure` necessitates the estimation of a *d*-dimensional linear subspace in ℝ^*n*^. The challenge is much more pronounced when the populations are highly admixed (large *σ*). Fig. 6 shows the model fits provided by `ALStructure`. Indeed, for large values of *σ* (*σ* = 2), `ALStructure` fails to correctly capture the admixture proportions. However, for smaller values of *σ* (*σ* = {1,0.5}), it can be seen that the fits provided by `ALStructure` are excellent. In all simulations *m* = 10^5^, *n* = 10^3^, and *d* = 3.

**Figure 6:**
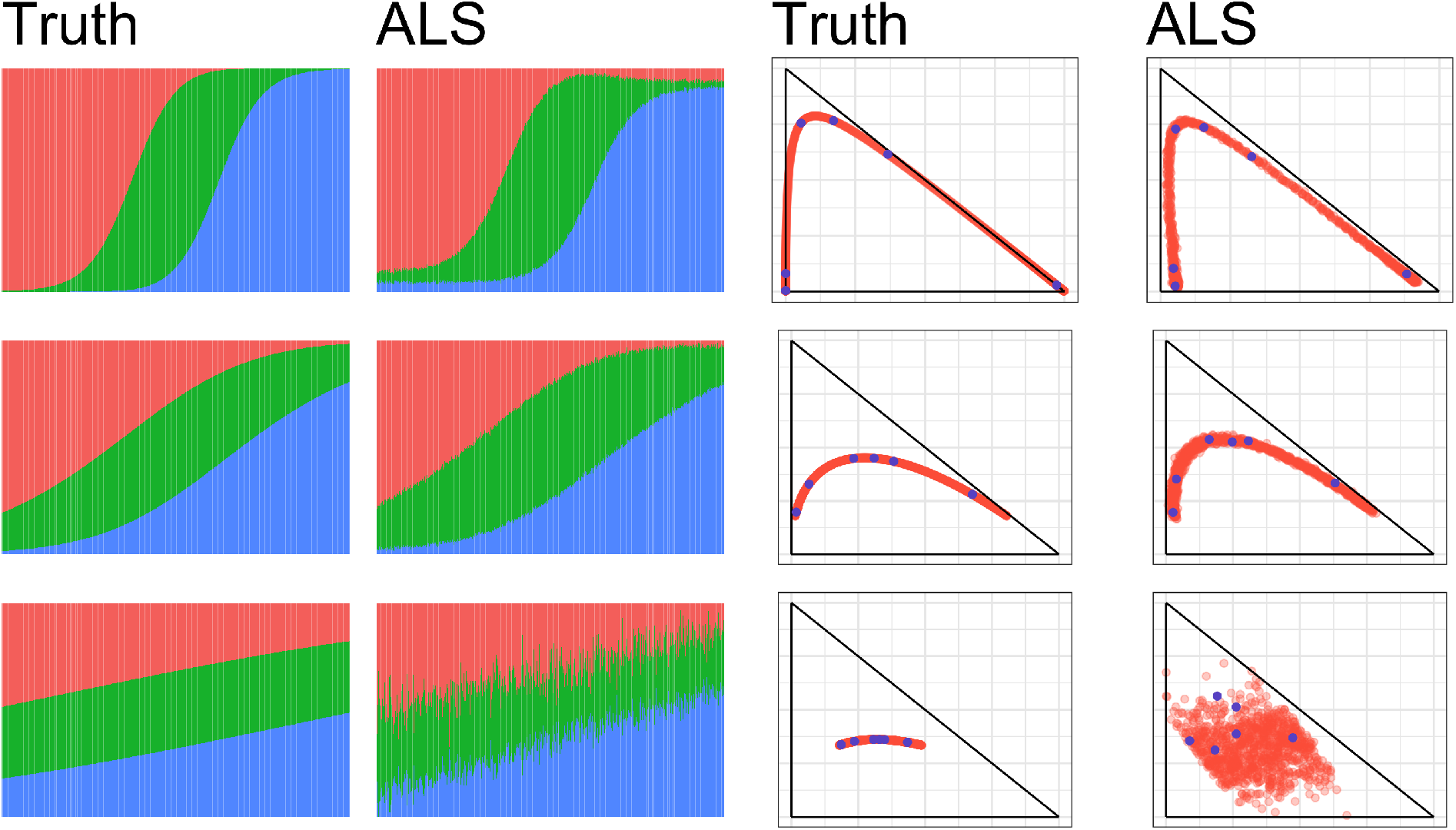
`ALStructure` fits of datasets from the Spatial model. (left) Stacked barplots of `ALStructure` fits. (right) Bi-plots of `ALStructure` fits. The parameter σ was set to 0.5, 1, and 2 for the top, middle and bottom rows, respectively. Blue points are plotted as a visual aid and delineate corresponding columns of ***Q*** and 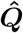.

We note that Gopalan *et al*. (2016) tested `Admixture`, `fastSTRUCTURE`, and `terastructure` on data drawn from the spatial model (which they refer to as “Scenario B”). They showed this model posed a significant challenge for all three methods, but found that `terastructure` performed the best.

## 5 Applications to global human studies

Here we apply `ALStructure` and existing methods to three globally sampled human genotype datasets: the Thousand Genomes Project (TGP), Human Genome Diversity Project (HGDP), and Human Origins (HO) datasets (Cavalli-Sforza, 2005; Lazaridis *et al*., 2014; The 1000 Genomes Project Consortium, 2015). Table 2 summarizes several basic parameters of each of the datasets and Appendix D details the procedures used for building each dataset. Although we recommend using sHWE from Hao and Storey (2017) for choosing *d*, here we take directly from Gopalan *et al*. (2016) the number of ancestral populations *d* so that our results are easily comparable to those.

**Table 2:** Dataset parameters.

| Dataset | $m$ | $n$ | $d$ | $m \times n$ |
| --- | --- | --- | --- | --- |
| TGP | 1229310 | 1815 | 8 | $\sim 2.2 \times 10^9$ |
| HGDP | 550303 | 940 | 10 | $\sim 5.2 \times 10^8$ |
| HO | 372446 | 2251 | 14 | $\sim 8.4 \times 10^8$ |

Fig. 7 shows scatterplots of the first two rows of 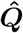 for each of the three datasets provided by each of the four fits. To disambiguate the inherent non-identifiability (see section 2.5), we ordered the rows of the fits 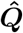 by decreasing variation explained: 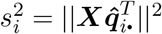. Perhaps the most striking aspect of Fig. 7 is the difference between the fits produced by each method. With the notable exception that `Admixture` and `ALStructure` have similar fits for the TGP and HGDP datasets, every pair of comparable scatterplots (i.e., within a single row of Fig. 7) are qualitatively different. Fig. 11 of Appendix F displays the same data represented as stacked barplots of the admixture proportions. In this representation too, qualitative differences between the fits are also evident. Table 3 shows the mean per-observation log-likelihood of the fits provided by each of the four methods. Supplementary Fig. 12 shows that the distributions of per-observation likelihood are nearly indistinguishable across all methods.

**Figure 7:**
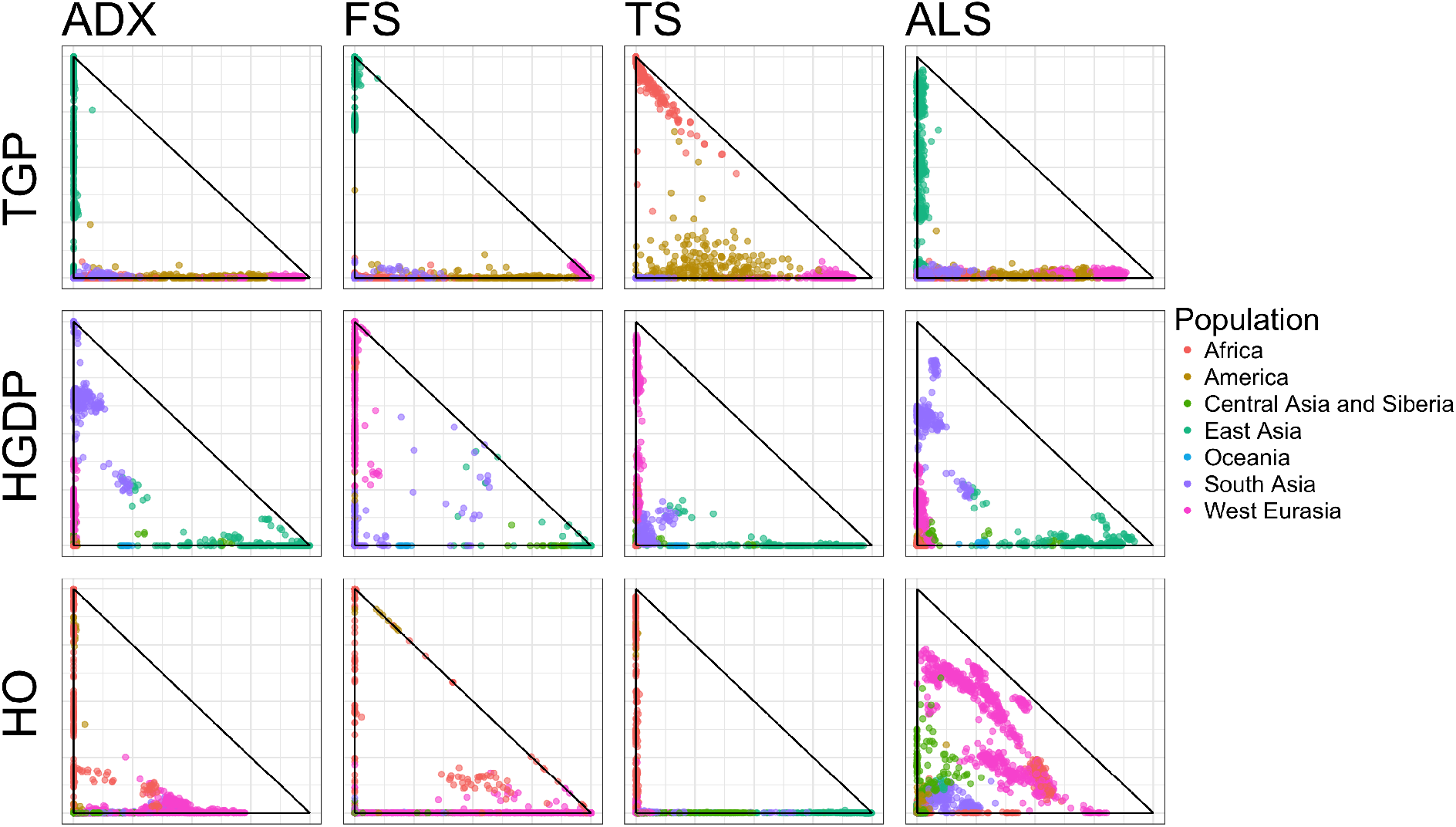
Bi-plots of the first two rows of ***Q*** (ranked by variation explained) of the fits of the TGP (top), HGDP (middle), and HO (bottom) datasets for each algorithm. Individuals are colored by coarse subpopulation from which they are sampled.

**Table 3:** The mean per-observation log-likelihood of each dataset under each method’s fit.

| Method | Admixture | fastSTRUCTURE | terastructure | ALStructure |
| --- | --- | --- | --- | --- |
| TGP | -0.7097 | -0.7136 | -0.7130 | -0.7100 |
| HGDP | -0.7494 | -0.7536 | -0.7608 | -0.7505 |
| HO | -0.7467 | -0.7515 | -0.7534 | -0.7477 |

Next we compare the performance of `ALStructure` to existing methods both in terms of efficiency and accuracy. Unlike in the case of simulated datasets where the ground truth is known, here we cannot directly compare the quality of model fits across methods. Instead, we assess the quality of each method by its performance on data simulated from real data fits. For concreteness, we briefly outline the process below:

(i) Fit each dataset with each of the four methods to obtain 12 model fits.
(ii) Simulate datasets from the admixture model using parameters obtained in the previous step.
(iii) Fit each of the 12 simulated datasets with each of the four datasets (48 fits) and compute error measures.

The process above treats each of the four methods symmetrically, evaluating each method based on its ability to fit data simulated from both its own model fits as well as every other methods’ model fits.

Fig. 8 summarizes the performance of each method with respect to both model fit and efficiency on data simulated from the above described process. As with the results on simulated datasets from Section 4.2, it is clear that `ALStructure` is competitive with respect to both model fit and time. Both `Admixture` and `ALStructure` outperform `fastSTRUCTURE` and terastructure by all quality of fit metrics. `ALStructure` far outperforms all methods with respect to time (one should note that the *y*-axis is on the *log* scale).

**Figure 8:**
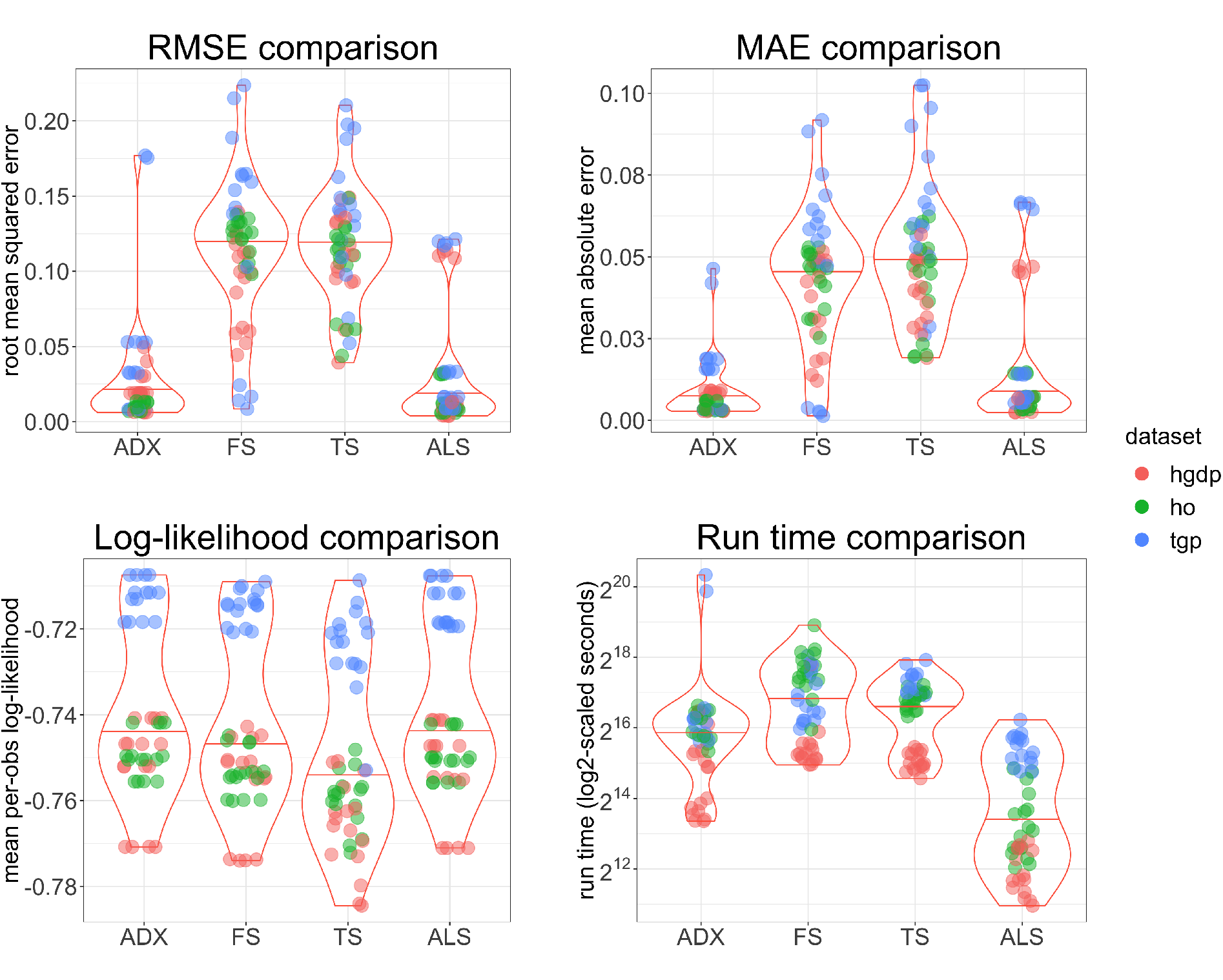
Summary of performance of `ALStructure` and preexisting algorithms on data simulated from real model fits. The points are colored by dataset.

In Appendix E we compare the performance of `ALStructure` to pre-existing methods on an additional non-global dataset from Basu *et al*. (2016). In this dataset, individuals are sampled from 18 modern Indian subpopulations. India’s genetic admixture is of particular interest because of its long history of sociocultural norms promoting endogamy. We find that each of the four methods produce admixture estimates qualitatively similar to each other for this dataset (see supplementary Figure 10). one possible explanation for this observed similarity is that the genetic history of India more closely mimics the admixture model than does global genetic history, as suggested by Lawson *et al*. (2018).

## 6 Discussion

In this work we have introduced `ALStructure`, a new method to fit the admixture model from observed genotypes. Our method attempts to find common ground between two previously distinct approaches to understanding genetic variation: likelihood-based approaches and PCA-based approaches. `ALStructure` features important merits from both. Like the likelihood-based approaches, `ALStructure` is grounded in the probabilistic admixture model and provides full estimates of global ancestry. However, operationally the `ALStructure` method closely resembles PCA-based approaches. In particular, `ALStructure`’s estimates of global ancestry are derived from a consistent PCA-based estimate that captures the underlying low-dimensional latent subspace. In this way, `ALStructure` can be considered a unification of likelihood-based and PCA-based methods.

Because `ALStructure` is operationally similar to PCA-based methods, it is computationally efficient. Specifically, the only computationally expensive operations required by the `ALStructure` algorithm are singular value and QR decompositions. Both of these computations have been extensively studied and optimized. Although `ALStructure` already performs favorably compared to preexisting algorithms in computational efficiency, it is likely that by applying more sophisticated matrix decomposition techniques `ALStructure` may see significant improvements in speed. Although extremely simple, `ALStructure` typically outperforms preexisting algorithms both in terms of accuracy and time. This observation holds under a wide array of datasets, both simulated and real.

The usefulness of PCA-based approaches has been increasingly recognized in related settings, such as the mixed membership stochastic block model (Rubin-Delanchy *et al*., 2017) and topic models (Ke and Wang, 2017). The basic approach we have presented is quite general. In particular, the set of models that satisfy the underlying assumptions of LSE is large, subsuming the admixture model as well as many other probabilistic models with low intrinsic dimensionality. Consequently, we expect that the `ALStructure` method can be trivially altered to apply to many similar problems beyond the estimation of global ancestry.

## Acknowledgements

Thank you to Wei Hao and Alejandro Ochoa for feedback and advice on this manuscript. This research was supported in part by NIH grant HG006448.

## Software

An R package implementing the method proposed here is available at https://github.com/StoreyLab/alstructure. Publicly available data sets were used in this study. The authors are able to provide the computers scripts utilized to analyze the data.

## Appendices

## A A Additional mathematical details

### A.1 Proof of Lemma 1

First we show that 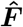 is unbiased. Note that:

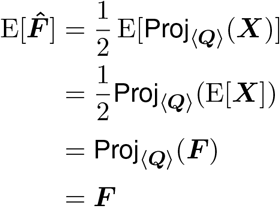

Between the first and second line, we note that the projection operator is linear and take advantage of linearity of expectation. Between the second and third line, we used the observation that 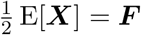. Finally, Proj_〈***Q***〉_ ***F*** = ***F*** since all rows of ***F*** belong to 〈***Q***〉. From an identical argument one can see that for projection onto any other subspace 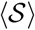, the corresponding estimator 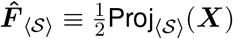 will have the property that

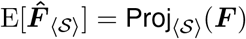

It is clear that if 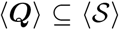, then 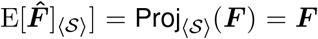 since the projection operator acts as the identity operator for vectors belonging to the subspace 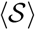.

Next we show that the converse is true: 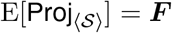 implies 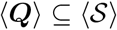. To do do this, we prove the contrapositive statement. If 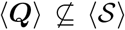 then 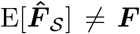. This can be seen by noting that each row in 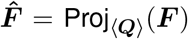 is a vector in the linear subspace 〈***Q***〉 projected into the linear subspace 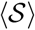; rows of 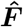 therefore belong to the linear subspace 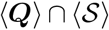. Unless 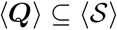, then the dimension of 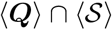 is strictly less than *d*, the dimension of 〈***Q***〉 and the rank of ***F***. Therefore, if 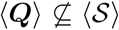, the rank of 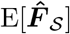 will be less than the rank of ***F***, implying 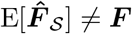.

### A.2 Proof of Lemma 2

Note that we can write the squared Frobenius norm as follows:

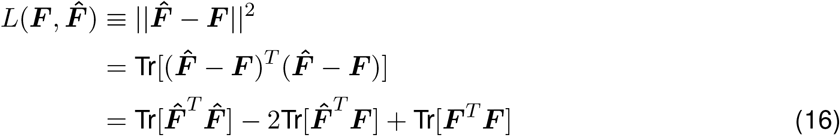

First let us compute the risk of our projection estimator 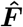. Suppose we have an orthonormal basis {***v***_*i*_} of 〈***Q***〉. Using the definition of 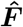 from Eq. 6 and the fact that the rows of both ***F*** belong to 〈***Q***〉, we note that we can write any row of either matrix in terms of the basis vectors {***v***_*i*_}:

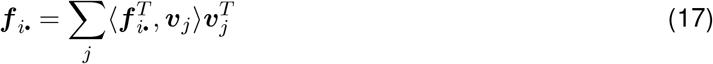

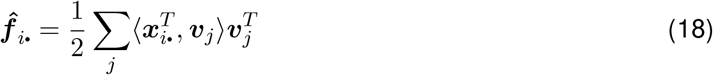

By rewriting the matrices 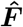 and ***F*** with respect to the basis {***v***_*i*_} and using Eqs. 17 and 18, it is a straightforward calculation to show that

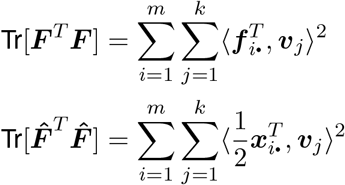

Substituting this result into Eq. 16 and taking expectations, we have the following expression for our loss function:

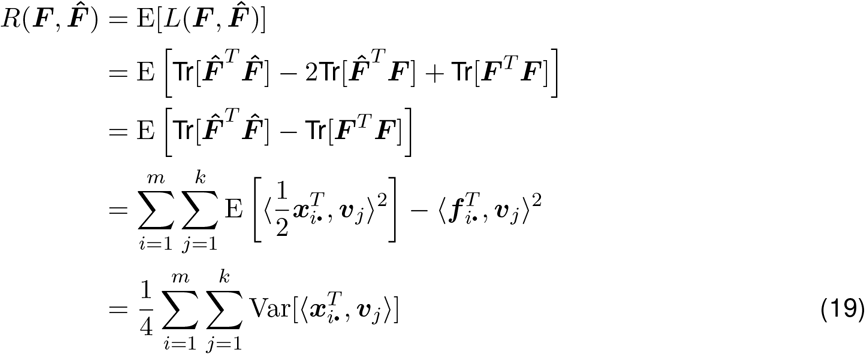

By studying Eq. 19, we can see the estimator 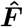 has several favorable properties. First note that the risk is a sum of *m × k* nonnegative numbers since Var[***Z***] ≥ 0 for any random variable ***Z***. If we were to project onto a larger subspace 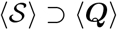, we would add terms to Eq. 19 and consequently increase our risk. If we were to project onto a smaller subspace 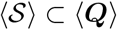, then the risk may decrease, however our new estimator will now be biased by Lemma 1. From these observations, we conclude that 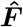 is optimal in the sense described in the Lemma 2.

### A.3 Proof of sufficiency of anchors

Here we show that either a set of anchor SNPs or a set of anchor individuals is sufficient to specify a unique factorization ***F*** = ***PQ*** up to the non-identifiability associated with row permutations.

#### Proposition 1.

*For a rank d matrix* ***F*** *with a factorization* ***F*** = ***PQ***, *if there is a set S of d rows of* ***P*** *such that for each i* ∈ {1,2,…,*d*} *there exists a row vector* ***p***_*i*._,. ∈ *S such that* ***p***_*i*._ = *δ_i_***e**_*i*_ *for δ* ≠ 0, *then the factorization is unique up to permutation. When such a set S exists, we say that we have “anchor SNPs.”*

*Proof*. Let us denote the matrix ***D*** = diag(*δ*_1_, *δ*_2_,…,*δ_d_*). Without loss of generality, let us assume that *S* is the first *d* rows of *P* and are ordered such that

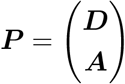

for some (*m* − *d*) × *d* matrix ***A***. Then there is a unique ***Q*** for this ***F*** matrix (up to permutation) which is

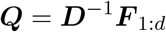

The matrix ***A*** is also uniquely determined by ***F*** once ***Q*** is fixed. To see this, note that

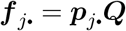

where ***f***_*j*._ and ***p***_*j*._ denote the *j* row of ***F*** and ***P*** respectively. Since ***f***_*j*._ is fixed and ***Q*** is unique under the anchor SNP assumption, there is a unique solution for ***p***_*j*._ by the linear independence of the rows of ***Q***.

The interpretation of the anchor SNPs assumption is that every ancestral population has at least one SNP that appears only in it. Presence of such an SNP is therefore a guarantee that the individual is a member of a particular population. Note that an identical argument could be made when we have a set *S* of *d* columns of ***Q*** that have exactly one nonzero entry at unique locations. When such a set exists, we say that we have “*anchor individuals*.” Under the admixture model, the simplex constraint requires that the nonzero entry of each anchor genotype is exactly one. In this scenario, there exists at least one individual from each ancestral population whose entire genome was inherited by a single ancestral population. We summarize these results in the following corollary and visualize the anchor SNP and anchor genotype scenarios in Fig. 1.

#### Corollary 1.

*Whenever a rank d matrix* ***F*** *admits a factorization* ***F*** = ***PQ*** *such that there are either a set of anchor SNPs or a set of anchor genotypes, the factorization is unique up to permutation*.

## B tALS and cALS comparisons

Fig 9 displays the 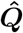 estimates of the `tALS` and `cALS` algorithms on simulated data from each of the four *α*-prototypes described in Section 4.1. For each of these datasets, *m* = 10^5^, *n* = 500, *d* = 3. We see that estimates provided by the `tALS` and `cALS` algorithms agree very well with each other for all *α*-prototypes. However, the run times are substantially different between these two methods, as displayed in Table 4: `tALS` terminates in minutes while `cALS` terminates in hours. Notably, the run times of the `tALS` algorithm also appear to be less sensitive to the *α*-prototype than the `cALS` algorithm. Most notably, the `cALS` algorithm takes an order of magnitude longer to run on the *α*_4_ prototype than any of the other *α*-prototypes. These observations support our preference for the `tALS` algorithm over the `cALS` algorithm.

Under *α*_2_ and *α*_3_, the estimates provided by `tALS` and `cALS` also agree very well with the true ***Q*** matrices. This is not the case under *α*_1_ and *α*_4_, where both algorithms provide substantially different results than the ground truth. However, because both of these *α*-prototypes lack a complete set of anchor SNPs, the model may well be unidentifiable.

**Table 4:** RMSE between true and estimated ***Q*** matrices for each method and each *α*-prototype (rows 1 and 2). RMSE between two estimated ***Q*** matrices (row 3). Run time (rows 4 and 5).

| | $\alpha_1$ | $\alpha_2$ | $\alpha_3$ | $\alpha_4$ |
| --- | --- | --- | --- | --- |
| $\text{RMSE}(Q, \hat{Q}_{\text{tALS}})$ | $6.2 \times 10^{-2}$ | $1.4 \times 10^{-2}$ | $8.7 \times 10^{-3}$ | $1.9 \times 10^{-1}$ |
| $\text{RMSE}(Q, \hat{Q}_{\text{cALS}})$ | $6.3 \times 10^{-2}$ | $1.5 \times 10^{-2}$ | $8.5 \times 10^{-3}$ | $2.2 \times 10^{-1}$ |
| $\text{RMSE}(\hat{Q}_{\text{tALS}}, \hat{Q}_{\text{cALS}})$ | $4.2 \times 10^{-3}$ | $2.3 \times 10^{-3}$ | $5.7 \times 10^{-4}$ | $3.9 \times 10^{-2}$ |
| tALS run time | 2.6 min | 1.4 min | 1.5 min | 2.8 min |
| cALS run time | 4.1 hr | 3.0 hr | 3.5 hr | 36.8 hr |

**Figure 9:**
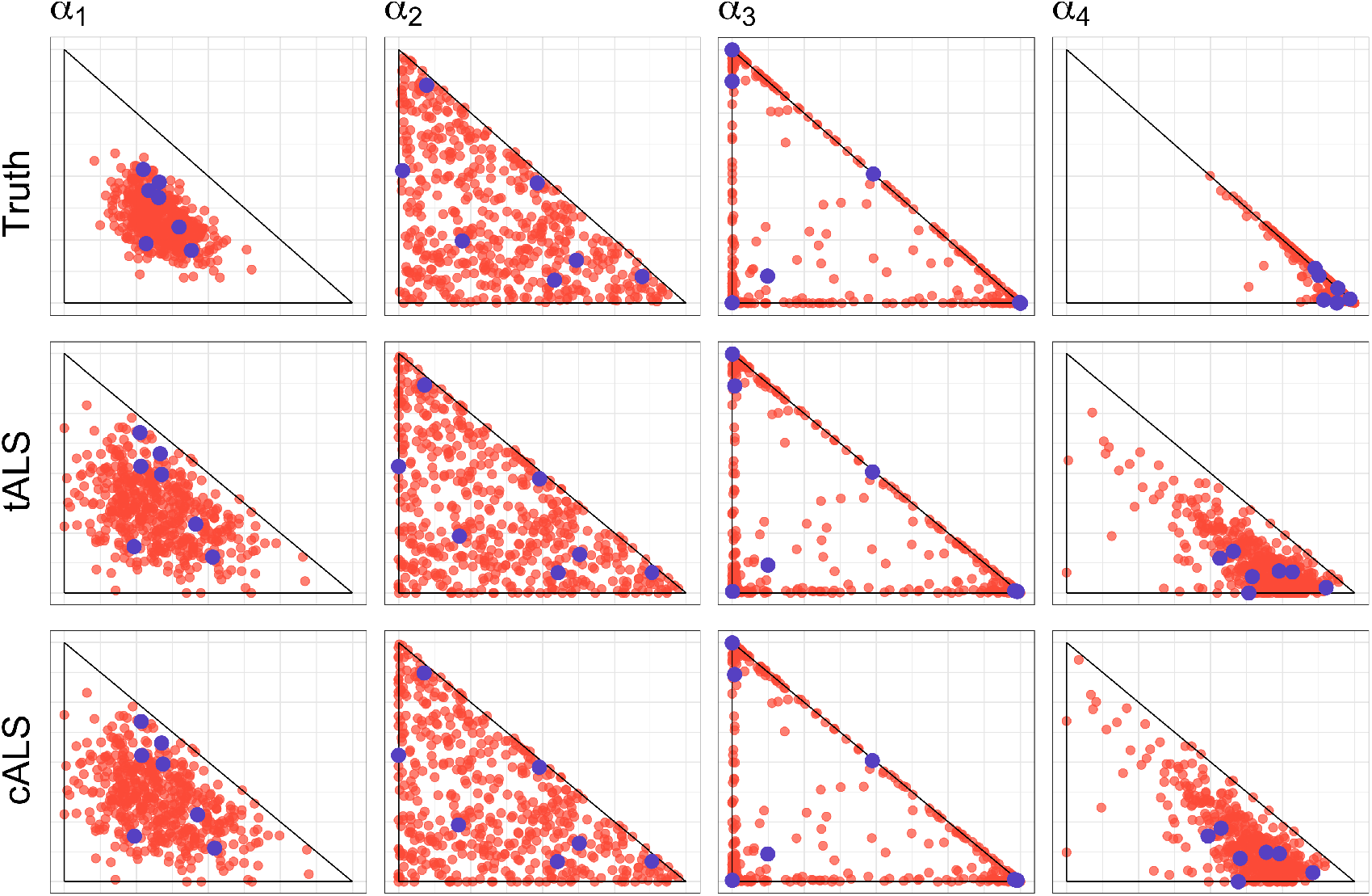
Biplots of the first two rows of ***Q*** (top), 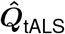 (middle) and 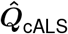 (bottom) for each of the four *α*-prototypes.

## C Simulation details

Due to time and computational constraints, each algorithm did not terminate on each of the 96 datasets generated for the simulations. In all, 326 of the 384 total simulations terminated during the one week time limit with a budget of 300 GB. `Admixture` completed 81 simulations, `fastSTRUCTURE` completed 80 simulations terastructure completed 81 simulations, and `ALStructure` completed 84 simulations. For the sake of comparison, Fig. 5 only shows the datasets for which all algorithms terminated.

## D HGDP, TGP, and HO dataset details

In Section 5 we analyze human genotype data from globally-sampled individuals. These data come from three public sources: HGDP (Cavalli-Sforza, 2005), TGP (The 1000 Genomes Project Consortium, 2015), and HO (Lazaridis *et al*., 2014). The various pre-processing steps are detailed below for each dataset.

### TGP

The 1000 Genomes Project dataset (TGP) samples globally from 26 populations and is available here: ftp://ftp.1000genomes.ebi.ac.uk/vol1/ftp/release/20130502/supporting/hd_genotype_chip/. Related individuals and SNPs with minor allele frequency < 5% are removed. The dimensions of this dataset are 1,716 individuals and 520,036 SNPs.

### HGDP

The Human Genome Diversity Project dataset (HGDP) samples globally from 51 populations and is available here: http://www.hagsc.org/hgdp/files.html. Individuals with first- or second-degree relatives and SNPs with minor allele frequency < 5% are removed. The dimensions of this dataset are 940 individuals and 550,303 SNPs.

### HO

The Affymetrix Human Origins dataset (HO) samples globally from 147 populations and is available here: http://genetics.med.harvard.edu/reich/Reich_Lab/Datasets.html. Nonhuman or ancient samples and SNPs with < 5% minor allele frequency are removed. The dimensions of this dataset are 2,248 individuals and 372,446 SNPs.

## E Application to a non-global dataset

In this appendix we apply `ALStructure` and preexisting methods to a dataset from Basu *et al*. (2016). In this dataset, individuals from 18 mainland Indian subpopulations are sampled. Following Basu *et al*. (2016), we set *d* = 4 for each method. Figure 10 plots the first two rows of ***Q*** output from `Admixture, fastSTRUCTURE, terastructure`, and `ALStructure`, respectively. As in the results from Section 5, rows of ***Q*** are ordered according to variation explained.

As can be seen, the estimated admixture proportions produced by each method are all qualitatively similar. Table 5 shows the likelihood of the data from each method, with each method performing similarly. The methods ranked by decreasing mean log-likelihood are: `Admixture, ALStructure, fastSTRUCTURE, terastructure`.

**Table 5:** Mean log-likelihood from of each method applied to Basu *et al*. (2016) dataset

| Method | Admixture | fastSTRUCTURE | terastructure | ALStructure |
| --- | --- | --- | --- | --- |
| Mean log-likelihood | -0.7360 | -0.7369 | -0.7373 | -0.7365 |

**Figure 10:**
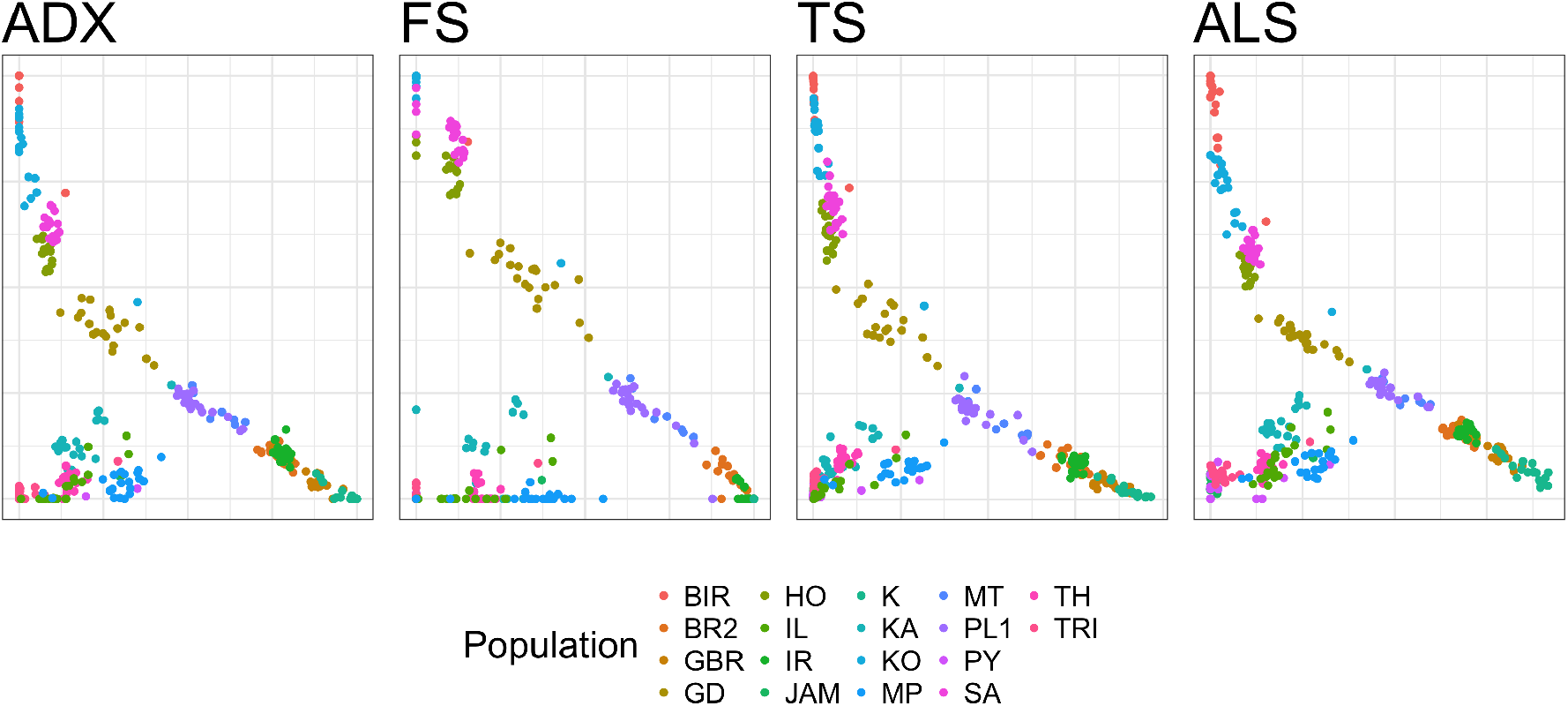
Bi-plots of the first two rows of ***Q*** (ranked by variation explained) of the fits of the Basu *et al*. (2016) dataset for each algorithm. Individuals are colored by the subpopulation from which they are sampled.

## F Supplementary figures

Figures start on the next page.

**Figure 11:**
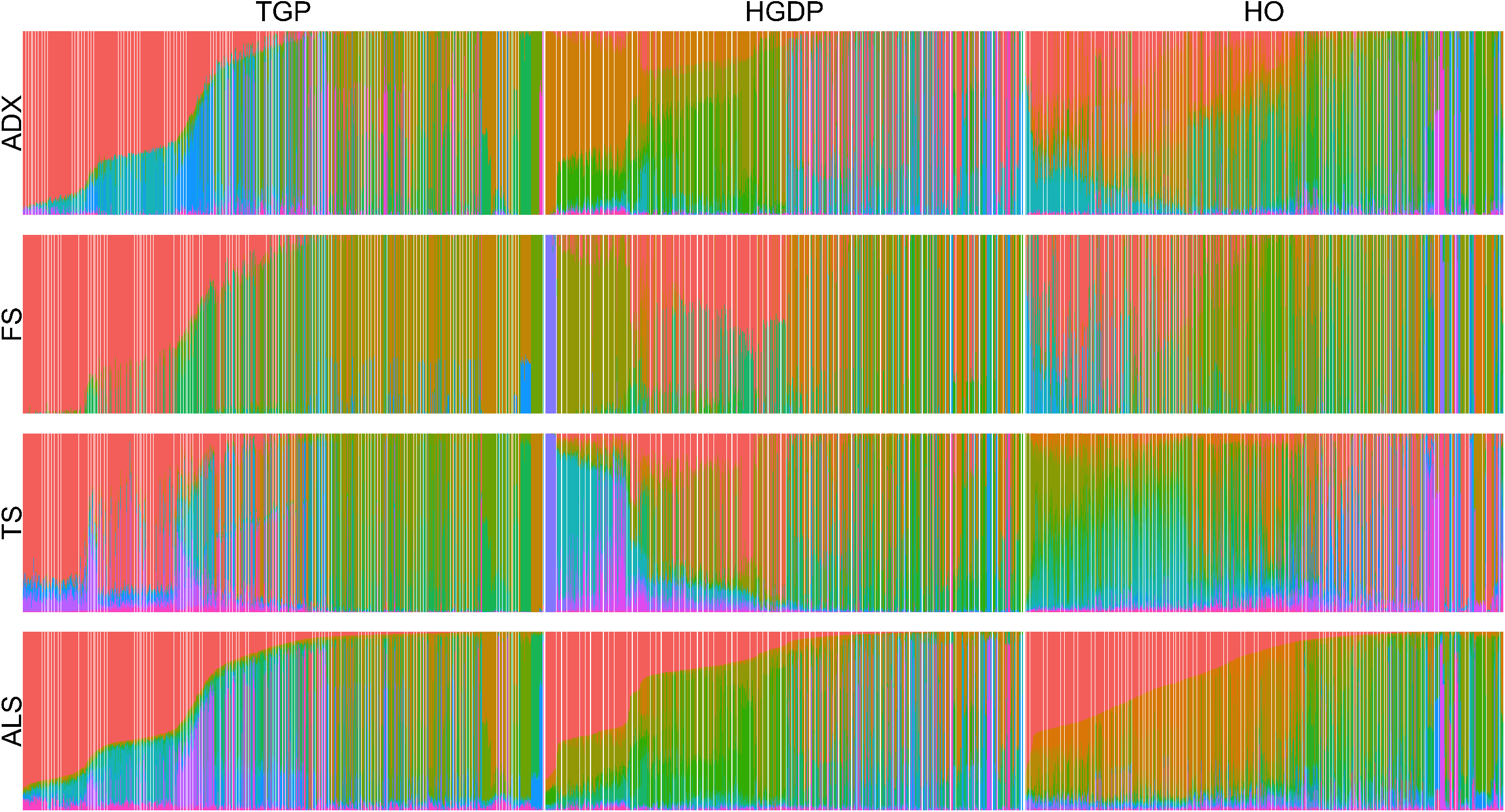
The admixture proportions of each globally sampled dataset as a stacked barplot. Because of the non-identifiability of the model, the order of the rows of ***Q*** are arbitrary. To disambiguate this, we order the rows of ***Q*** in each dataset by decreasing average admixture. The coloring in Fig. 11 is then done according to this ordering. As an aid to the eye, we also reorder the columns of ***Q*** according to decreasing proportion of the first row of ***Q*** of the `ALStructure` fit. The choice to order all fits according to `ALStructure` is arbitrary, however all fits must be ordered consistently to make meaningful comparisons possible. As can be seen, each of the fits differ significantly from each other on every dataset.

**Figure 12:**
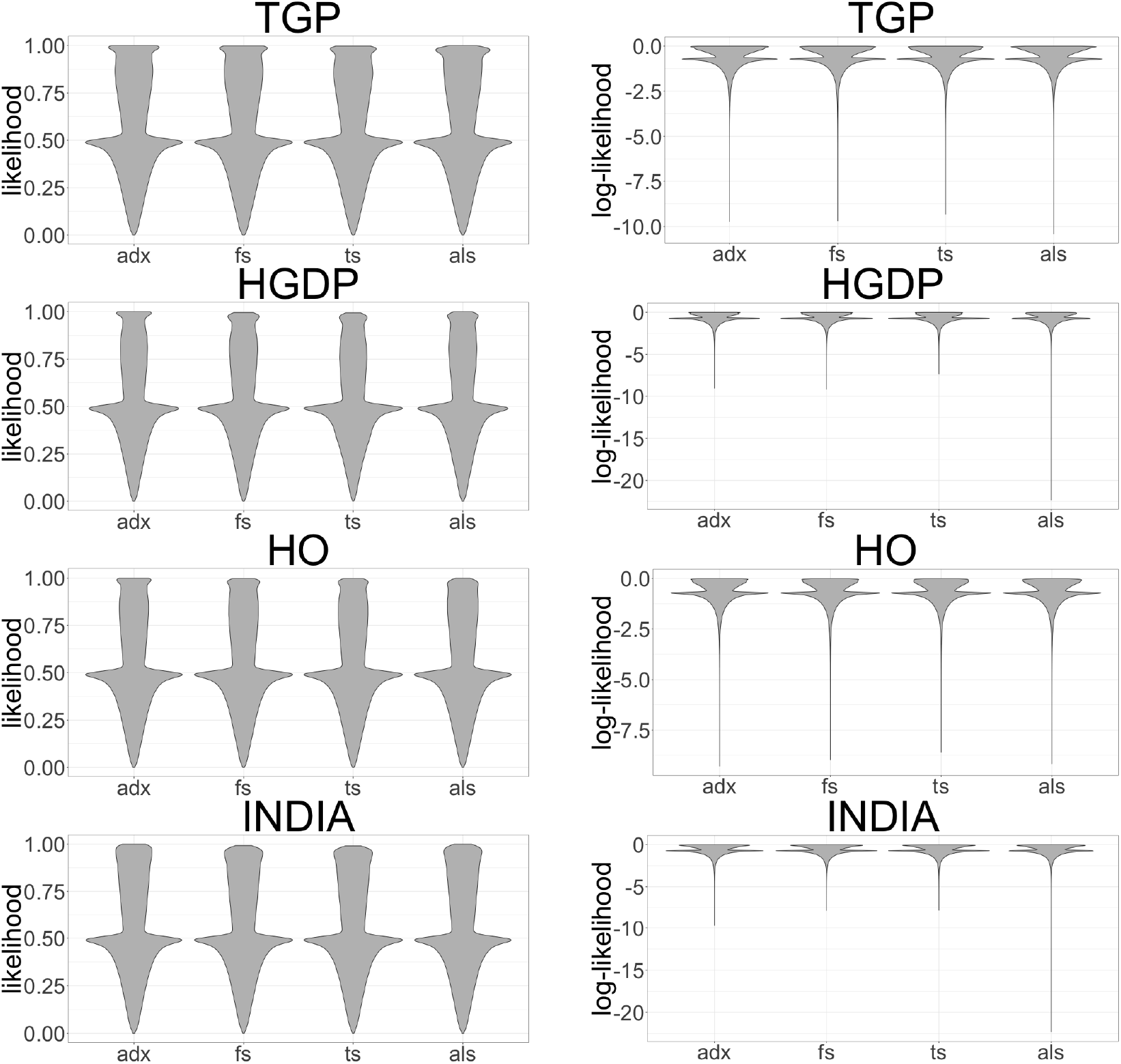
(left) The distribution of likelihoods for each element of ***X*** for each method and dataset. (right) The same as (left) except on a log-scale.

## Footnotes

1 Although Tipping and Bishop (1999) introduced a probabilistic interpretation of PCA for multivariate normal data, to our knowledge no such interpretation of PCA exists when the data are binomial, as is the case in the admixture model.

2 Other prior distributions can be used for ***P*** and ***Q*** (Pritchard *et al*., 2000), but here we refer to the PSD model as that using the priors listed here.

3 Refer to Chen and Storey (2015) for a precise statement of the theoretical assumptions of LSE.

4 The notation 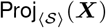 typically refers to projection of the columns of ***X*** onto the linear subspace 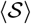, but here we use this notation to denote projection of the rows of ***X*** onto 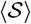.

5 The result from Grippo and Sciandrone (2000) is actually more general than this. We reproduce the special case above in order to make clear the connection to our problem.

6 We note that we have decided to use the `tALS` function rather than the `cALS` function in our definition of the `ALStructure` algorithm, valuing the speed advantage of `tALS` over the theoretical guarantees of `cALS`. If desired, one could of course choose to use the `cALS` function instead without making any other alterations to the `ALStructure`.

